# Deep Mining of the Human Antibody Repertoire Identifies Frequent and Genetically Diverse CDRH3 Topologies Targetable by Vaccination

**DOI:** 10.1101/2024.10.04.616739

**Authors:** Rumi Habib, Shahlo O. Solieva, Zi Jie Lin, Sukanya Ghosh, Kelly Bayruns, Maya Singh, Colby J. Agostino, Nicholas J. Tursi, Kirsten J. Sowers, Jinwei Huang, Ryan S. Roark, Mansi Purwar, Younghoon Park, Kasirajan Ayyanathan, Hui Li, John W. Carey, Amber Kim, Joyce Park, Madison E. McCanna, Ashwin N. Skelly, Neethu Chokkalingam, Sinja Kriete, Nicholas Shupin, Alana Huynh, Susanne Walker, Roopak Sadeesh, Niklas Laenger, Jianqiu Du, Jiayan Cui, Ami Patel, Amelia Escolano, Peter D. Kwong, Lawrence Shapiro, Gregory R. Bowman, Beatrice H. Hahn, George M. Shaw, David B. Weiner, Jesper Pallesen, Daniel W. Kulp

## Abstract

Germline targeting vaccination strategies against highly variable pathogens such as HIV aim to elicit broadly neutralizing antibodies (bnAbs) with particular immunogenetic or structural features. The V2 apex of the HIV Env protein is a promising target for a class of bnAbs that contain conserved structural motifs in the heavy chain complementarity determining region 3 (CDRH3). Here, we show that these structural motifs are targetable by vaccination by characterizing V2 apex ‘axe-like’ CDRH3s in the human repertoire and developing new immunogens capable of engaging them. We determined the frequency and diversity of axe-like CDHR3s in healthy human donors using a series of structural informatics approaches, finding these precursors in nearly 90% of donors. Axe-targeting immunogens based on the HIV Env Q23.17 bound axe-like precursors in cryo-EM structures, induced V2 apex-specific antibody responses in humanized mice, and induced axe-like heterologous neutralizing antibodies in rhesus macaques infected with a germline-targeted simian-human immunodeficiency virus. These results illustrate a new structure-guided immunoinformatic vaccine design paradigm that can be employed to elicit immunogenetically diverse yet structurally conserved classes of antibodies.

**Significance Statement:** Many broadly neutralizing antibodies (bnAbs) utilize modes of epitope recognition dominated by the antibody complementarity determining region 3 (CDRH3). The CDRH3 is the most diverse part of the antibody, posing a challenge for germline targeting vaccine designs that aim to elicit antibodies with particular immunogenetic features. Vaccine design strategies that accommodate CDRH3 variability are therefore needed. Many HIV Env V2 apex bnAbs share “axe-like” CDRH3 microfolds that arise from diverse immunogenetic origins. Here we determined the frequency in humans of B cells with such CDRH3 topologies and designed immunogens to engage their precursors. This work opens a path toward vaccines that engage specific structural classes of B cells, thereby advancing the rational design of immunogens for HIV and other pathogens.

## Introduction

Following the identification of SARS-CoV-2, a highly efficacious vaccine for the virus was developed in less than a year. Yet, despite over 35 years of research, there is still no protective vaccine for HIV. The HIV envelope glycoprotein (Env), which is the only target for neutralizing antibodies, has several features that make it a challenge for vaccine development. These include a large and dynamic glycan shield, conformational masking of epitopes, and extensive sequence diversity (1–4). As a result of these properties, the antibody response to Env is usually strain-specific targeting surface-exposed hypervariable sequence motifs (5). However, 10-20% of infected individuals develop broadly neutralizing antibodies (bnAbs) that target conserved but recessed epitopes on Env (6–12). These antibodies have been shown to be protective against infection by diverse heterologous viruses in humans and rhesus macaques (13–16). Consequently, the reproducible elicitation of bnAbs is a primary goal of most HIV vaccine designs.

Germline targeting approaches to bnAb elicitation focus on identifying key features encoded by the germline immunoglobulin gene repertoire critical for epitope engagement by prototype bnAbs and aim to engineer immunogens capable of engaging and expanding those germline precursors (1, 8, 17–20). Among the various classes of HIV bnAbs, antibodies targeting the V2 apex are some of the most frequently elicited during infection, are highly potent, and have relatively straightforward developmental pathways (21–24). These features make the apex a promising target for vaccine development. V2 apex bnAbs primarily rely on their long heavy chain third complementarity determining regions (CDRH3s) for binding to Env, forming distinct microdomains that contact several glycans and the C-strand of the V1V2 region (23, 25–34). These CDRH3s can be roughly classified into three categories based on their structural topology: “needle-like” in which β-hairpin CDRH3s bearing sulfated tyrosines at their tips extend into the trimer apex hole (23, 25, 26); “axe” or “hammerhead-like” in which CDRH3s form extended β-sheet interactions with the C-strand (27–30); and “combined,” which share features of both (31, 32). Because of these unusual structural features and uncommon length (>21 aa, IMGT numbering), B cells expressing V2 apex bnAb precursors are estimated to have low frequencies in the naïve human B cell repertoire, making priming a major hurdle to their elicitation (24, 25). This challenge is compounded by the fact that, unlike bnAb precursors to other epitopes, V2 apex bnAbs utilize diverse V, D, and J genes, making it difficult to target V2 apex bnAb precursors based on sequence features alone (24, 25, 32). Yet, despite their diverse gene usage, V2 apex bnAbs form reproducible structural classes (27, 32). In line with this, immunogenetically diverse antibodies to the V2 apex have been isolated from rhesus macaques that recapitulate the V2 apex bnAb microdomains found in humans, (23, 32–35) indicating that a myriad of genetically diverse CDRH3s can converge on structurally convergent solutions to engage the V2 apex (32). Here, we highlight that V2 apex bnAb precursors can be identified by the structural topology of their CDRH3s, and that by targeting structural rather than immunogenetic classes of antibodies, it may be possible to capture a greater diversity and broaden the opportunities for engaging rare bnAb precursors (Fig. 1A).

**Figure 1.**
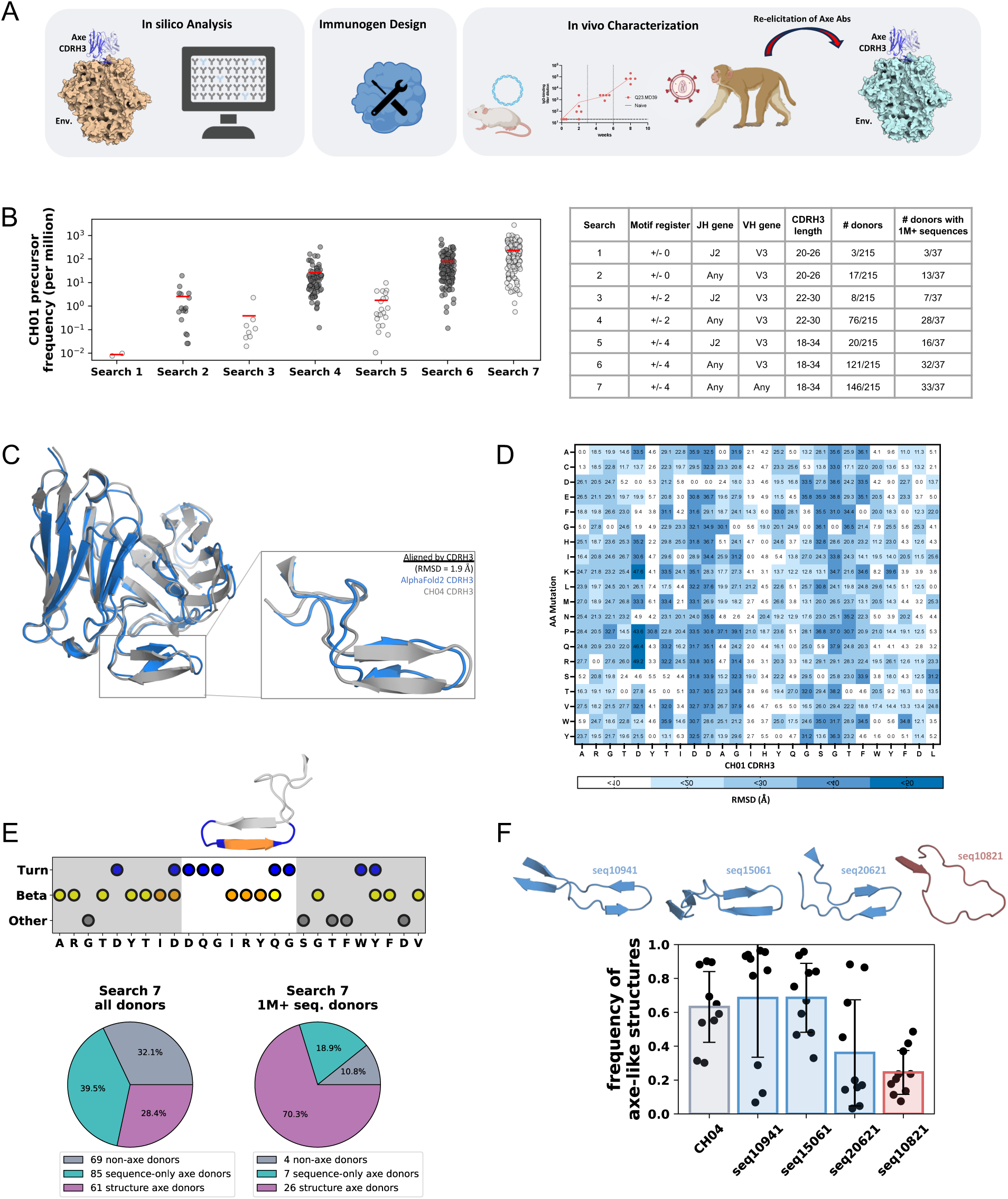
Using structural characteristics of axe-like bnAbs to estimate axe-like B cell frequency in the human repertoire. **A.** An overview of our study, starting from *in silico* analysis to immunogen design to *in vivo* characterization and re-elicitation of axe-like antibodies in rhesus macaques. **B.** CH01-like antibody search results from the OAS database. Each point on the graph represents a donor that had at least one sequence matching the search criteria. All search criteria used are detailed in the table (right). The DH gene motif found in the CH01 lineage, Y[YQK]GSG, was used for the searches, however the searches did not filter by a specific DH gene. **C.** An AlphaFold2 model of CH01 (blue) has a low CDRH3 Cα RMSD to the published CH04 structure (gray, PDB ID: 3TCL). **D.** AlphaFold models from *in silico* CDRH3 saturation mutagenesis where each position along the CH01 CDHR3 was mutated to every possible residue. The x-axis shows the native CH01 CDRH3 sequence while the y-axis shows the mutations tested. The minimum CDRH3 Cα RMSD (Å) to the AlphaFold-generated control structure is shown for each mutation. Small RMSD values (white, light blue) represent positions where the mutations did not greatly affect the fold while large RMSD values (darker blue) represent positions where the mutations greatly perturbed the CDRH3 fold. **E.** The scatter plot shows our structural definition of an axe CDRH3 fold applied to CH04 (PDB ID: 3TCL), where blue shows residues that are involved in a turn, yellow shows beta-sheet residues, orange shows beta-sheet residues that are at least 20 Å away from the start of the CDRH3, and gray shows residues that did not fit our turn or beta criteria. The pie chart on the left shows the distribution of donors (n=215) that have no CH01-like B cells (gray), donors that have at least one CH01-like B cell sequence but was not predicted to have any axe-like structures (cyan), and donors that have at least one CH01-like B cell sequence predicted to fold into an axe-like structure. The pie chart on the right conducts the same analysis on donors with at least 1M sequences in the OAS database (n=37). **F.** Structures of antibodies found from Search 7, three of which were predicted to be axe-like (blue) and 1 which was predicted to not be axe-like (red). The plot below the structures shows the ratio of frames within each MD simulation where the structure was classified as axe-like. Error bars represent one standard deviation from the mean.

Axe-like V2 apex bnAb lineages are typified by the human bnAb lineages PG9 and CH01 (36, 37). To understand the structural plasticity and frequency of axe-like bnAb lineage precursor B cells, we first conducted large-scale sequence and structure prediction screens to identify the frequency of CH01-like, PG9-like, and rhesus axe-like antibodies in the human B cell repertoire, identifying this class of antibodies as a promising target for vaccination. Next, we designed and characterized a stable recombinant Env trimer immunogen, Q23.MD39, based on the clade A Env Q23.17, which exhibits affinity for the inferred germline precursors (iGLs) of the PG9 and CH01 lineages (27, 36, 38, 39). Because we had previously shown that immunization with DNA-encoded native-like HIV trimers can enhance antibody responses compared to conventional protein bolus vaccination in WT mice (40), we first delivered Q23.MD39 using DNA immunization and showed that it can assemble into well-formed trimers *in vivo*. To further enhance the antigenicity of this construct, we next engineered self-assembling ferritin nanoparticles decorated with Q23.MD39. These nanoparticles showed enhanced engagement of axe-like iGLs and could be delivered by DNA vaccination to assemble *in vivo*. We took advantage of the high affinity of Q23.MD39 for the CH01 iGL to solve an atomic resolution structure of this antibody, finding that it retains the same unique axe-like CDRH3 conformation as the mature CH01 antibody. We further enhanced the affinity of Q23.MD39 for additional axe-like iGLs through a single-round mammalian display method and showed that it can elicit axe-like neutralizing antibodies to the V2 apex as determined by cryo-EM in SHIV-infected macaques and V2 apex-targeted responses in humanized mouse models.

## Results

### Estimation of axe-like B cell frequency in the human repertoire

Currently, four mature axe-like V2 apex bnAb lineages have been described. These include the rhesus lineages V033-a and 41328-a (32), as well as the prototypical human lineages CH01 (36) and PG9 (28). The V033-a, CH01, and PG9 bnAb lineages are promising examples of V2 apex bNAbs given their high breadth, potency, and relatively low levels of somatic hypermutation. Even though the CH01 lineage has the second-shortest CDRH3 length of canonical human V2 apex bnAbs, previous estimates of B cell precursor frequency have suggested that CH01-like B cells may be one of the rarer classes of V2 apex bnAbs (25). This was primarily due to CH01’s rare D-J gene pairing and its skewed D gene placement in its CDRH3. For the CH01 bnAb lineage, the major contribution of the D gene to the CDRH3 of the antibody is a “YYGSG” sequence motif that forms the C-terminal section of the CDRH3 axe. We hypothesized that CDRH3-dominated bnAb precursor searches should take into account two previously unconsidered criteria: 1) that antibodies with alternative CDRH3 lengths, J genes, and D gene placement could represent a larger pool of potential bnAb precursors, and 2) that only a subset of these sequences can form the requisite CDRH3 shape present in the target antibody class. Recent advancements in B cell receptor repertoire sequencing, structure prediction tools, and computational power make it possible to apply these criteria to identify potential bnAb precursors through this new structure informatic approach.

The Observed Antibody Space (OAS) database contains over one billion antibody sequences, allowing for an in-depth antibody repertoire analysis at a previously unachievable level (41). Furthermore, it is possible to model the antibody sequences within the OAS database with protein structure prediction software such as AlphaFold2 or AlphaFold3 (AF2/AF3) (42, 43). We conducted sequence searches within the OAS database to estimate frequencies of potential bnAb precursors. Our first set of searches was focused on the CH01 bnAb lineage. We conducted a precursor search of OAS imposing the following CH01 criteria: CDHR3 length of 26 amino acids, V3 gene family, J2 gene and D gene amino acids in the same location in the CDRH3 (“motif register”), similar to previously published criteria (25). These CH01 CDRH3 criteria resulted in an average frequency estimation of about 0.01 precursor per million B cells (‘Search 1’ in Fig. 1B). As the JH gene amino acids do not make direct Env contacts, we removed the JH gene restriction and observed 1 precursor per million B cells, and an increase in the number of donors with sequence matches (‘Search 2’ in Fig. 1B). This drastic increase relative to a previous study (25) was expected since the J2 gene utilized by CH01 is the least frequently used JH gene in the human B cell repertoire (44). The frequency then increased significantly to ∼100 precursors per million B cells when the CDRH3 length (20-40 residues in length), D gene motif position (+/- 2 to 4 residues) and V gene restrictions were relaxed (‘Search 3-7’ in Fig. 1B). Using the most relaxed search criteria, ‘Search 7’, we observed a frequency of 228 precursors per million B cells encoding a diverse set of 48,713 non-redundant antibody sequences. We further observed CH01-like sequences in 33 out of the 37 donors who have >1 million sequences in OAS (Extended Figure 1a). Taken together, these results suggest that CH01-like bnAb precursors may be more frequent in humans than previously estimated.

To test whether the sequences found from our OAS searches are predicted to have an axe-shaped CDRH3 fold we modeled them with AF2. First, we determined that AF2 could predict the known axe-like CDRH3 of CH01 with high structural precision (< 2Å RMSD) (Fig. 1C). To assess which mutations in CH01 were required for AF2 to predict the axe-like structure, we conducted an *in silico* saturation mutagenesis screen by predicting each point mutation within CH01’s CDRH3. Interestingly we observed positions D105, D106, and to a lesser extent positions G113, S114, G115, led to high structural deviation when mutated, suggesting these amino acids played a key role in this axe-like micro-fold of the CH01 CDRH3 (Fig. 1D). To better evaluate the ability of AF2 to predict the axe-like CDRH3 topologies on an unbiased set of antibodies, we predicted the structures of recently reported rhesus V2 apex bnAbs (32). Remarkably, AlphaFold2 was able to differentiate the needle- and axe-like CDRH3s of these mAbs, which were not included in the AlphaFold2 training set, and are immunogenetically distinct from published human V2 apex bnAbs. In some cases, we found that AlphaFold was able to generate predictions strikingly similar to published Cryo-EM structures (Fig. S2). While AF2 can struggle to model most CDRH3 structures (45–47), our data suggest that it can reliably predict the axe-like CDRH3 structure present in many V2 apex bnAbs.

Next, we sought to discover which of the BCR sequences from search criteria 7 could fold into axe-like structures with AF2. To automate detection of axe-like structures, we developed a general axe-like CDRH3 feature pattern that we termed ‘turn-beta-turn’ from axe harboring antibodies with known structures (Fig. 1E, Fig. S2A). The ‘turn-beta-turn’ pattern consists of at least two consecutive ‘turn’ residues defined using canonical turn metrics and types (48, 49), followed by two or more residues with ‘beta’ backbone geometry at least 20Å away from the start of the CDRH3 and capped by at least two more consecutive ‘turn’ residues (see methods). We validated the specificity of this ‘turn-beta-turn’ pattern by searching 610 diverse antibody structures which target many different proteins from the SAbDab database (50) and found five known axe-like structures harboring a ‘turn-beta-turn’ pattern (CH01 lineage, PDB IDs 3TCL, 3U46, 3U4B, and CAP256-VRC26 lineage antibodies, PDB IDs: 4ORD, 4ORG), and only three other antibodies which exhibit an axe-like fold (Fig. S2). Of the search 7 sequences, we found 5,146 of the sequences have axe-like structures. We found that, out of the donors with >1 million antibody sequences, 10.8% do not have any axe-like sequence or structure shaped antibodies, 18.9% have axe-like sequence signatures without an axe shape and 78.4% have a predicted CH01-like axe shaped CDRH3. To further support AF2 predictions, we used molecular dynamics (MD) simulations to assess the stability of the axe microfolds in our structural models (51). We conducted 10 replicate MD simulations of 250 ns each (aggregate of 2.5 ms per antibody) for four AF2 models and CH04 (PDB ID: 3TCL) as a control. We applied our axe structural definition to the simulations to determine the frequency of axe-like structures within each simulation. We found that the CDRH3 maintained an axe-like structure for a majority of the time in the CH04 simulations (Fig. 1F). Three antibodies that matched the ‘turn-beta-turn’ structural motif (seq10941, seq15061, seq20621, shown in blue in Fig. 1F) also maintained an axe-like structure in a majority of the simulations. The antibody that did not match the ‘turn-beta-turn’ structural motif (seq10821, shown in red in Fig. 1F) had a drastically lower axe-like frequency compared to CH04. These simulations gave us novel insights into the dynamics of the CDRH3 loops and showed that this approach can be a useful tool for verifying the stability of CDRH3s with predicted micro-folds, such as the axe shape.

The axe-like PG9 CDRH3 has a sequence signature that differs from CH01, and we sought to understand how frequently the PG9 sequence and structure could be found in human repertoires. We employed our expanded sequence and structure modeling pipeline on PG9 as we did for CH01 (Fig. S1B) and found PG9-like sequences in 35 out of 37 deeply sequenced donors (>1M sequences) (Fig. S1B). From the 38,427 PG9-like sequences, we found that 6,965 sequences had axe-like structures and that 81% of deeply sequenced donors have axe-like antibodies. These searches show that, similarly to CH01, the human antibody repertoire contains an immunogenetically diverse set of PG9-like antibodies (Fig. S1C).

### Rhesus V2 apex bnAb motifs are frequent in the human repertoire

The discovery of rhesus-derived axe-like bnAb lineages (V033-a and 41238-a) gives us additional targets for vaccine design (32). Interestingly, although human V2 apex bnAbs use diverse V, D, and J genes, a striking feature of rhesus V2 apex bnAbs is their nearly uniform usage of the IGHD3-15*01 gene (24, 32–35). These bnAbs invariably incorporate a D gene-templated “EDDYG” motif that is absent from D genes in the human repertoire. Therefore, an outstanding question is how frequently human V-D-J rearranged antibodies, which encode non-templated N-additions on either side of the D gene, contain this key “rhesus” sequence motif. To explore this, we conducted repertoire and structural searches to understand how often an “EDDYG” motif is incorporated into human CDRH3s.

We performed three searches using either a larger rhesus D gene sequence of “EDDYGYYT”, the core rhesus D gene motif of “EDDYG”, and the motif found in the V033-a lineage “[EG]DDYG” with two or more N-terminal amino acids and six or more C-terminal residues relative to the motif (Fig. 2A, Fig. S1D). We observed two human donors who harbored antibodies with the larger rhesus D gene sequence (Fig. 2A, Search 1). We identified CDRH3s in donors Hu-CD1 and Hu-BD3 (52) with striking similarity to rhesus lineages V033-a and 41328-a, suggesting a subset of the human population may have rhesus-like V2 apex bnAb precursors (Fig. 2B). While the full D-gene motif was rare amongst donors, the core D gene motifs were found in 89% and 97% of donors (Fig. 2A, Search 2 and Search 3, respectively). For CDRH3 lengths of >23, the core D-gene motifs were found at 6 per million and 8 per million antibody sequences. To further probe the immunogenetics of antibodies with these core D-gene motifs, we cataloged the human D-genes used in antibodies found in Search 2. We observed that IGHD4-11*01, IGHD4-17*01 and IGHD4/OR15-4a*01 each contained the ‘DYG’ sequence (Fig. 2B) and were all heavily enriched in the core D-gene search relative the frequency of these D genes found across OAS (Fig. 2C). We folded Search 3 CDRH3s using AF2 and found that 267 sequences out of the 24,696 sequences form axe-like structures (three axe-like and one non-axe examples are shown in Fig. 2D). Indeed, MD simulations of rhesus-like axe CDRH3s demonstrate that this fold is highly stable (Fig. 2E). Therefore, despite the nearly universal usage of an IGHD3-15*01 D gene-derived “EDDYG” motif in rhesus V2 apex bnAbs, the absence of an orthologous human D gene did not limit the generation of EDDY-like motifs in rearranged human V-D-J recombinants. Thus, V2 apex bnAb induction in the rhesus model may find parallels in human infection and vaccination.

**Figure 2.**
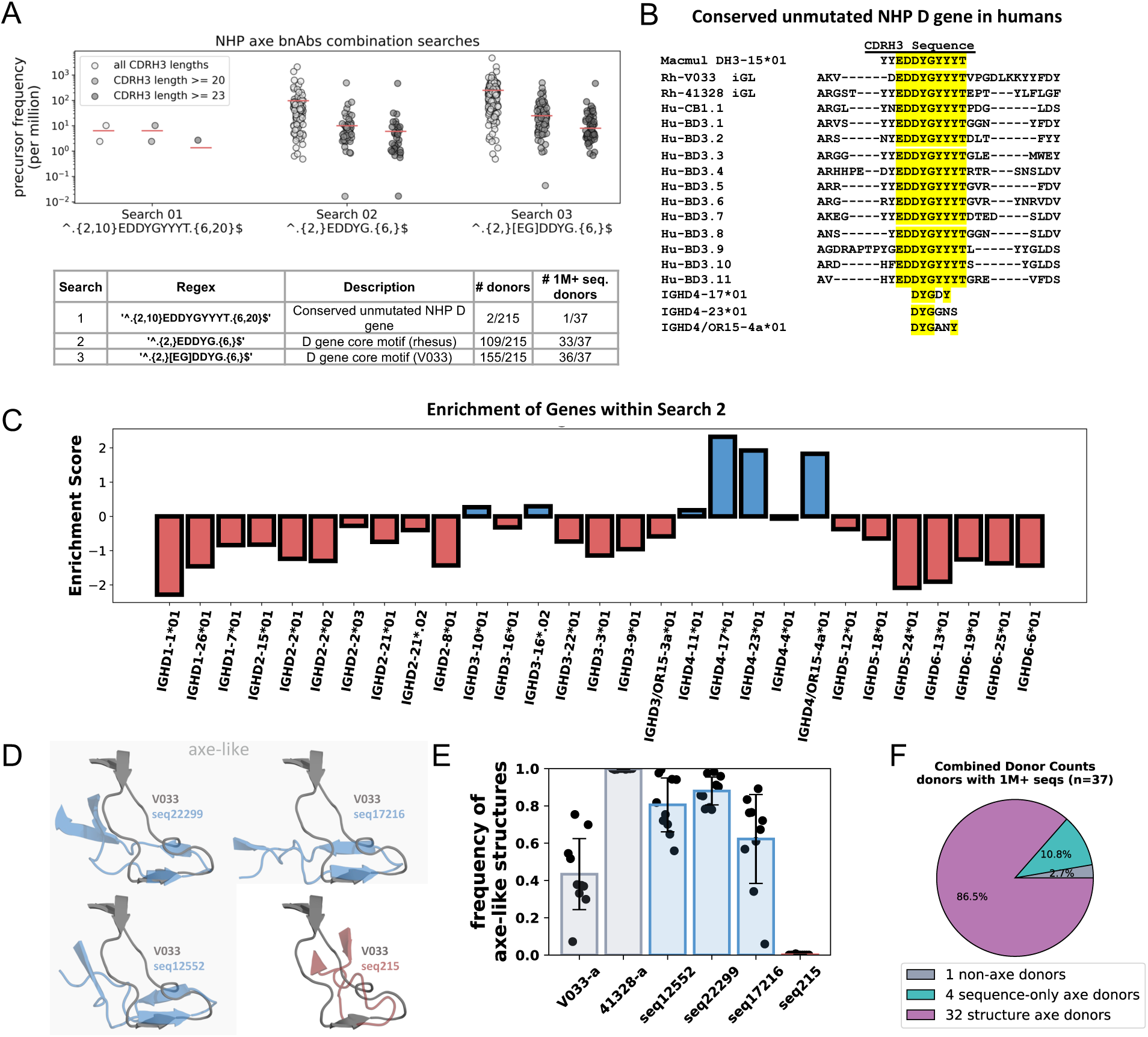
Estimating the frequency rhesus-like axe antibodies within the human repertoire. **A.** Various rhesus-like antibody search results from the OAS database, broken down into CDRH3 lengths (all lengths, at least 20 amino acids in length, and at least 23 amino acids in length). Each point on the graph represents a donor that had at least one sequence matching the search criteria. The three searches are detailed in the table. **B.** Sequence alignments of the antibodies from macaques V033 and 41328, rhesus and human D genes, and sequences found in Search 1. **C.** The enrichment of IGHD genes within Search 2, with positive enrichment in blue and negative enrichment in red. **D.** Structures of antibodies found in Search 3, 3 of which are classified as axe-like and 1 is not classified as axe-like. **E.** The ratio of frames within each trajectory where the structure was classified as axe-like. Error bars represent one standard deviation from the mean. **F.** Distribution of highly sequenced donors (1M+ sequences, n=37) that have no axe-like B cells (gray), donors that have at least one axe-like B cell sequence but was not predicted to have any axe-like structures (cyan), and donors that have at least one axe-like B cell sequence predicted to fold into an axe-like structure. Axe-like B cells are defined to include CH01-like, PG9-like, or rhesus-like sequences motifs shown in Fig. 1B (search 7), Fig. S1B (Search 7), and Fig. 2A (Seach 3), respectively.

While only four mature axe-like V2 apex bnAbs have thus far been described, it is likely that far more exist with these sequence and structure features. Using the four axe-like bnAbs as a reference, we found 12,378 total axe-like antibody sequences. Of the donors with > 1 million antibody sequences in OAS, we found that 1 donor did not have any axe-like CDRH3s, but 97% of donors had antibodies with axe-like *sequence* motifs and 86% of donors had antibodies with axe-like *structural* motifs (Fig. 2F). Collectively, these studies reveal that B cells with axe-like sequence motifs are identified at high frequency in the human antibody repertoire and are observed in the overwhelming majority of donors, pointing to this structural class of bnAbs as a promising vaccine target.

### Design, characterization, and nucleic acid delivery of a Q23.17-based native-like trimer and nanoparticle immunogens

To design an immunogen that can engage axe-like V2 apex bnAb precursors, we sought to develop a stable trimeric Env protein that could easily multimerize on nanoparticles and be delivered by nucleic acid platforms while retaining its structure *in vivo*. To discover a starting immunogen sequence, we searched for Envs that have detectable affinity for inferred germline precursors of V2 apex bnAbs, as we reasoned that these may be more favorable for V2 apex priming. Several groups have conduct screens of Envs to identify such strains with “permissive” V2 epitopes that have natural propensities for V2 apex iGL engagement (27, 36, 38). We selected a clade A Env, Q23.17 (53), as our platform for immunogen development because of its relatively high binding affinity for the iGLs of the axe-like CH01 and PG9 bnAb lineages (27, 36, 38), sensitivity to mature V2 apex bnAbs, and frequent elicitation of axe-like V2 apex bnAbs in SHIV-infected rhesus macaques (24, 32). We note that this Env also contains many of the engineered “Apex-GT5” residues recently reported to sensitize the Env BG505 to precursors of the PCT64 lineage (25, 33).

To design a stabilized recombinant Q23.17 trimer, we tested several published Env trimer stabilization strategies including MD39 (54), MD64 (55), v5 (56), 7S (57), RnS-DS (58, 59), and Olio6 (55) mutations. We found that Q23.MD39 and Q23.7S demonstrated superior expression and antigenicity. The MD39 mutations are a subset of the 7S mutations, and thus we concluded that the MD39 mutations were primarily responsible for the stable phenotype of the trimer and focused on Q23.MD39 for further investigation. Q23.MD39 bound strongly by ELISA to the trimer-specific or trimer-preferring mAbs PGDM1400, PGT151, and VRC34 and bound poorly to V2p-, CD4i-, and linear V3-specific non-neutralizing antibodies, indicating an antigenic profile consistent with a stable prefusion trimer (Fig. 3A). In contrast, Q23.gp120.foldon, a construct consisting of three gp120 domains loosely held together by a foldon trimerization domain, showed high binding to non-nAbs and poor binding to quaternary structure-dependent bnAbs (Fig. 3A). We confirmed the size and homogeneity of Q23.MD39 by size exclusion chromatography multiple angle light scattering (SEC-MALS) which showed that Q23.MD39 formed trimers of the expected size (Fig. S3B).

**Figure 3.**
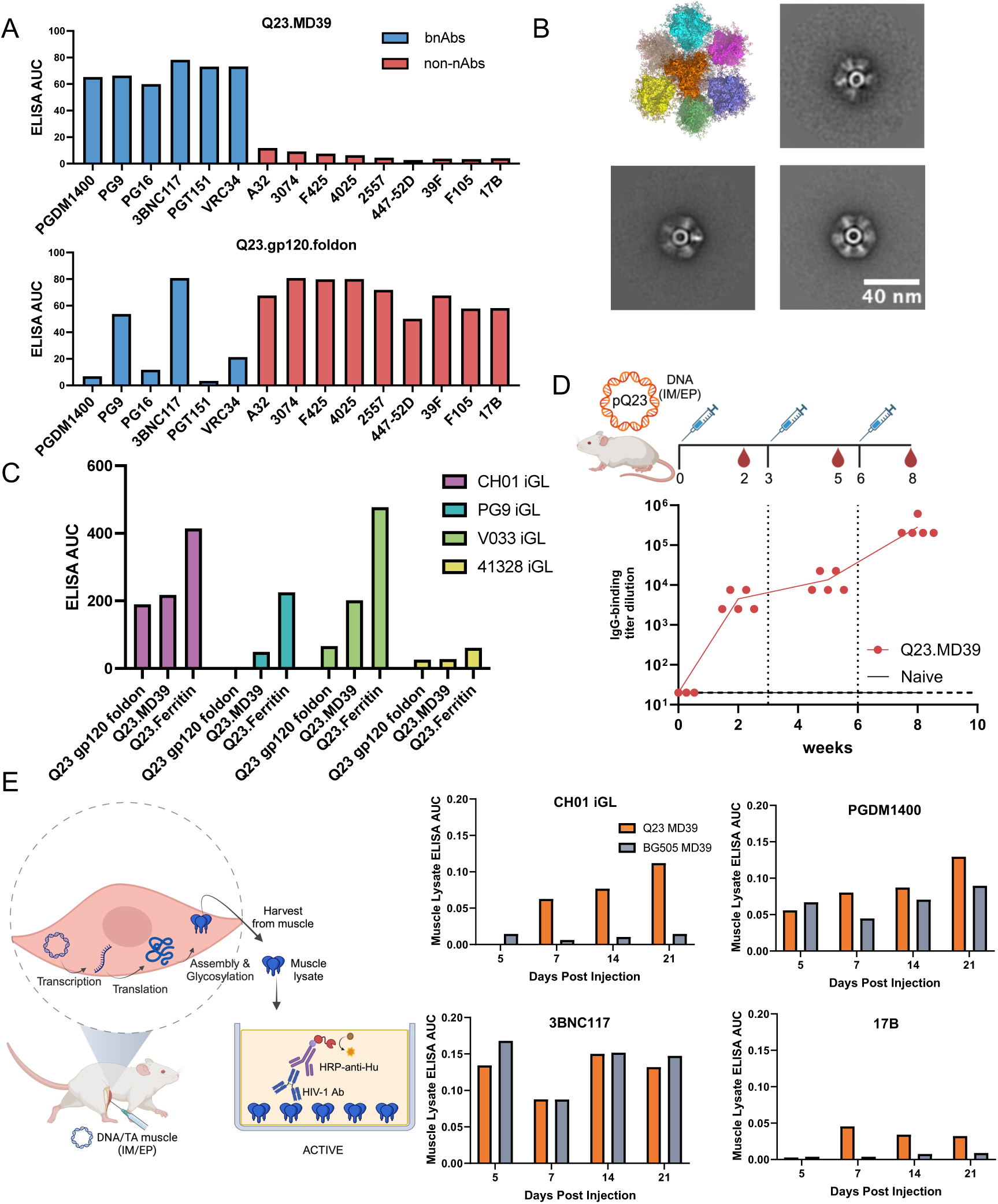
Characterization and DNA-delivery of Q23-based immunogens. **A.** antigenic profile of Q23.MD39 and Q23.gp120.foldon against a panel of broadly neutralizing and non-neutralizing antibodies. **B.** Model and NS-EM 2D class averages of Q23.Ferritin nanoparticles. **C.** ELISA area under the curve (AUC) binding of Q23.gp120.foldon, Q23.MD39, and Q23.Ferritin to axe-like V2 apex bnAb iGLs. **D.** Time course of vaccine-specific serum-IgG binding endpoint titers in WT BALB/c mice immunized with 25µg of DNA-delivered Q23.MD39. Circles indicate individual mice, and vertical dotted lines indicate immunization timepoints. e, Left, overview of the ACTIV assay. Right, ELISA area under the curve binding of muscle lysate from mice immunized with Q23.MD39 (orange) or BG505.MD39 (grey) to various antibodies.

There is extensive evidence that the multimeric display of vaccine antigens can significantly enhance their immunogenicity through increased avidity, enhanced trafficking, or other effects (60–64). This result has been observed for V2 apex immunogens particularly (33, 34, 65). We have previously shown that DNA-launched nanoparticles (DLNPs) are capable of self-assembly *in vivo* and are highly immunogenic (66). Thus, we engineered Q23.Ferritin, a self-assembling ferritin nanoparticle decorated with our Q23.MD39 trimer (Fig. S7). Through NS-EM and antigenic profiling by ELISA, we determined that this nanoparticle retained a closed, prefusion phenotype and expressed primarily as a well-formed nanoparticle based on SEC analysis (Fig. 3B, Fig. S3C). We measured the binding of our design in soluble trimer, nanoparticle, and gp120 forms to the CH01, PG9, V033, and 41328 iGLs, which demonstrated that Q23-based immunogens have affinity for axe-like iGL precursor antibodies (Fig. 3C).

DNA delivery of HIV Env trimers is a promising approach for driving neutralizing antibody responses in WT mice (40). Importantly, SEC analysis showed that Q23.MD39 expressed primarily as a trimer (Fig. S3A). Unlike purification of recombinant Env immunogens, nucleic acid delivery strategies are unable to exclude poorly folded or unassembled Env proteins. To show that the trimer was capable of homogenous assembly and expression *in vivo*, we immunized WT BALB/c mice with a DNA vaccine encoding the Q23.MD39 construct. Mice were immunized at weeks 0, 3 and 6, with 5ug, 10ug or 25ug of DNA encoded Q23.MD39 with or without an IL-12 genetic adjuvant, and antibody responses were tracked for 2 weeks following each immunization (Fig. 3D, Fig. S4A). The immunized animals developed strong binding antibody responses, demonstrating that the trimer was immunogenic when delivered as a DNA vaccine (Fig. 3D, Fig. S4A). However, they failed to develop neutralizing antibody responses or V2 apex-targeted responses. To determine if the trimer structure and concomitantly the antigenicity of Q23.MD39 produced *in vivo* was maintained, we employed our recently developed Antigen Conformation Tracing In Vivo by ELISA (ACTIVE) assay (40) (Fig. 3E). Mice were immunized with DNA containing Q23.MD39 or BG505.MD39 intramuscularly. *In vivo* assembled trimers from harvested muscle tissue of mice were analyzed by ELISA for binding to various bnAbs, non-nAbs, and the CH01 iGL at several timepoints post-administration (Fig. S4B). High binding of muscle homogenate expressing Q23.MD39 to trimer-specific bnAbs such as PGDM1400, and low binding to non-nAbs such as 17B, indicated that Q23.MD39 retains its favorable antigenic profile when expressed *in vivo* (Fig. 3E, Fig. S4B). BG505.MD39, a similarly stabilized and closely related Env, showed a similar binding profile to Q23.MD39. However, only Q23.MD39 was able to bind to CH01 iGL *in vivo* (Fig. 3E, Fig. S4B). Escalating-dose immunization regimes delivered over the course of a few weeks have been shown to significantly improve immune responses through an increase in durable germinal center responses (67, 68). Here, with a single DNA immunization, we detected increasing amounts of intact prefusion Env trimer in muscle tissue for at least 3 weeks post-immunization (Fig. 3E, Fig. S4B). These data thus show that Q23.MD39 retains its sensitivity for V2 bnAb iGLs *in vivo*.

We next sought to understand whether Q23.Ferritin could also be delivered by DNA vaccines. We have previously demonstrated that DNA immunizations can be used to deliver self-assembling nanoparticle immunogens such as eOD-GT8 60mer (66), and others have shown that Env trimer nanoparticles can be delivered by mRNA (69). The favorable expression profile of Q23.Ferritin combined with our use of a genetic fusion approach again allowed us to deliver the immunogen via DNA immunization (Fig. S3C). DNA delivery of the Q23.Ferritin construct with or without plasmid IL-12 in WT mice elicited high-titer binding antibody responses, demonstrating that Q23.Ferritin is immunogenic when delivered by DNA vaccination (Fig. S4A).

### Cryo-EM structure of Q23.MD39 bound to the CH01 iGL reveals key molecular contacts of axe-like precursor antibodies

To gain structural insights into how Q23.MD39 binds to the axe-like CH01 iGL with high affinity, we determined atomic resolution cryo-EM structures of Q23.MD39 in complex with the Env gp120-gp41 interface bnAb 35O22 alone (3.19Å), in complex with the CH01 iGL and 35O22 (3.35Å), or in complex with the mature CH01 bnAb and 35O22 (3.49Å) (Fig. 4A, Table S1). We observed a prefusion-closed Q23.MD39 Env trimer with high structural similarity to BG505.SOSIP.664, further indicating that Q23.MD39 is a well-formed native-like trimer. Overlay of our CH01 iGL and mature antibody structures showed identical conformations for all CDR loops except the CDRL1 (Fig 4B). The CH01 iGL antibody bound to Q23.MD39 Env trimer with a 1:1 stoichiometry (Fig. 4A) and engaged the C-strand using an intricate backbone hydrogen bond network similar to that found in our structure of the mature CH01 bnAb, as well as in previous structures of the lineage members CH03/CH04 in complex with a C-strand scaffolded protein (Fig. 4C, top left panel) (27). The sidechains of the C-strand on protomer A interacted with positions within the axe shape: Y100g with R169 and Y100h with K168 (Fig. 4C, top middle and top right panels). The sidechains of the CDRH1 interacted with both the K171 and Y173 on the gp120A C-strand (38) (Fig. 4C bottom left panel). The sidechains on the CDRH2 and CDRH3 formed extensive contacts with the N160A glycan (Fig. 4C bottom middle panel). The structure of the CH01 iGL maintained the axe-like CDRH3 of the mature CH01-lineage bnAbs, suggesting that this unique CDRH3 topology was likely a feature present in the unmutated ancestor of the bnAb lineage. This finding corroborates the structure-guided immunoinformatic approach to bnAb precursor identification conducted in this study, as it suggests that CH01-like germline precursors also harbor this unique CDRH3 shape.

**Figure 4.**
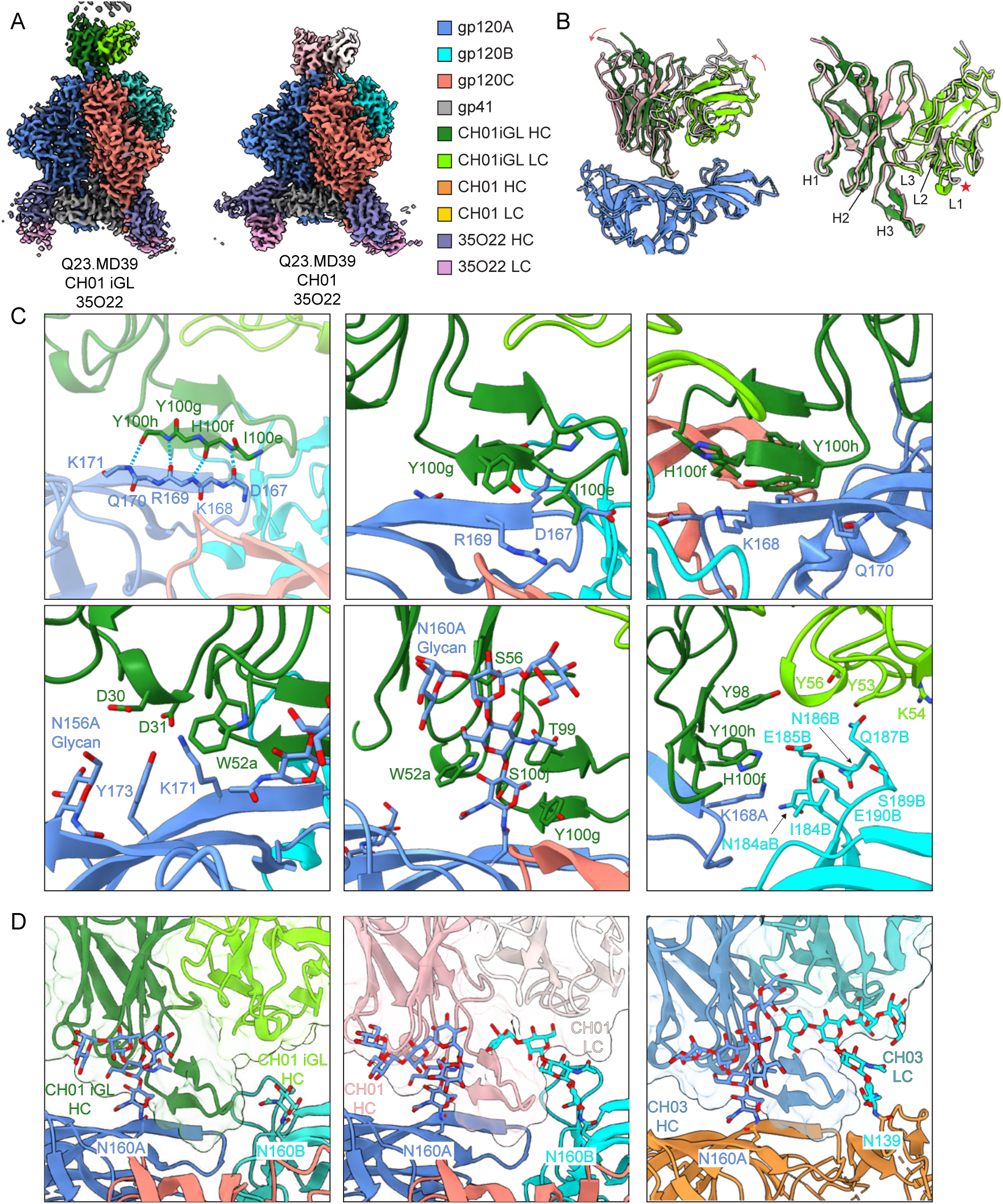
Cryo-EM structure of Q23.MD39 in complex with the CH01 iGL. **A.** Colored density maps of structures of Q23.MD39 in complex with 35O22 alone (left) or CH01 iGL and 35O22 (right). **B.** Overlayed atomic models of CH01 iGL, CH03 (PDB: 5ESV), and CH04 (PDB: 5ESZ). **C.** Interactions at the epitope-paratope interface of CH01 iGL and Q23.MD39. **D.** Glycan interactions between the CH01 iGL with N160 (left), the mature CH01 bnAb with N160 (middle), and the CH03 bnAb with N160 and N139 (PDB: 5ESV).

Upon comparing our CH01 iGL structure with the mature CH01 structure, we found the two models overlapped perfectly at the CDRH3 loop but that the mature CH01 bnAb Fab had tilted slightly further towards the Env protomer interacting with the heavy chain. Notable features that differed from the CH01 iGL in the mature bnAb structure were a more pronounced electron density for the N156A and N160B glycan trees and adjustment of the light chain to move further away from the V2b hypervariable loop (Fig 4D). The more pronounced electron density and closer contact with the N156 and N160 glycans indicated that the mature antibody had affinity matured towards accommodation of these conserved V2 apex glycans. Interestingly, the modeled N160B glycan in our CH01-bound Q23.MD39 structure occupied the same groove that was occupied by the N139 glycan tree in the previously solved CH03 crystal structure. Furthermore, the adjustment of the light chain to move further away from the V2b hypervariable loop indicates that CH01 affinity matured to avoid this site of high variability. Together, these features of the mature CH01 bnAb likely contribute to its improved breadth and potency compared to the CH01 iGL.

From this structural analysis, we hypothesized there are three major factors contributing to the high affinity of Q23.17 for the axe-like CH01 iGL: (i) C-strand residues that are optimal for interactions with the CH01 iGL CDRH3; (ii) a lack of disfavored glycans at the apex epitope such as N130 and loop V2b glycans as previously reported (38); and (iii) the length of loop V2b being within the threshold accommodated by the CH01 iGL, in particular by the CDRL2 and CDRH3. This observation suggests V2b could be a promising location for incorporating designed loops or mutations, with the closest contact in our structure being CH01 iGL HC Y100h and gp120B E185 (4.82 Å). With this structure, we can now visualize an axe-like precursor antibody engaging with the C-strand and rationally design potential mutations that may increase affinity for CH01-like axe-shaped precursors.

### Mammalian display mutagenesis to enhance axe-like iGL binding

While Q23.MD39 has modest binding affinity for known axe-like iGLs, the iGLs available for axe-like human bnAb lineages contain affinity matured non-templated regions in CDRH3. Since these contribute a significant portion of the epitope-paratope interface, authentic naïve B cell unmutated common ancestor (UCA) receptors are likely to exhibit substantially lower affinities for Env. Therefore, we hypothesized that further optimization of Env is likely required to elicit true axe-like antibodies with high efficiency. To this end, we used a novel mammalian display mutagenesis approach to enhance the affinity of Q23.MD39 for axe-like iGLs. We optimized a previous mammalian display protocol (54), by performing single round sorting followed by next-generation sequencing. Briefly, Q23.MD39 was membrane anchored using a PDGFR transmembrane domain and a lentiviral scanning NNK library spanning Env residues 110-192 was created with silent barcoding mutations flanking each NNK codon. The top 5% of cells were sorted against axe-like iGLs and deep sequenced (Fig. 5A). To identify affinity-enhancing mutations, the frequency of particular mutants in the sorted population was compared to the frequency of the same mutation in unsorted cells. This method enabled the identification of a significantly higher number of affinity-enhancing mutations with greater efficiency and speed compared to traditional mammalian display mutagenesis approaches (Fig. 5B).

**Figure 5.**
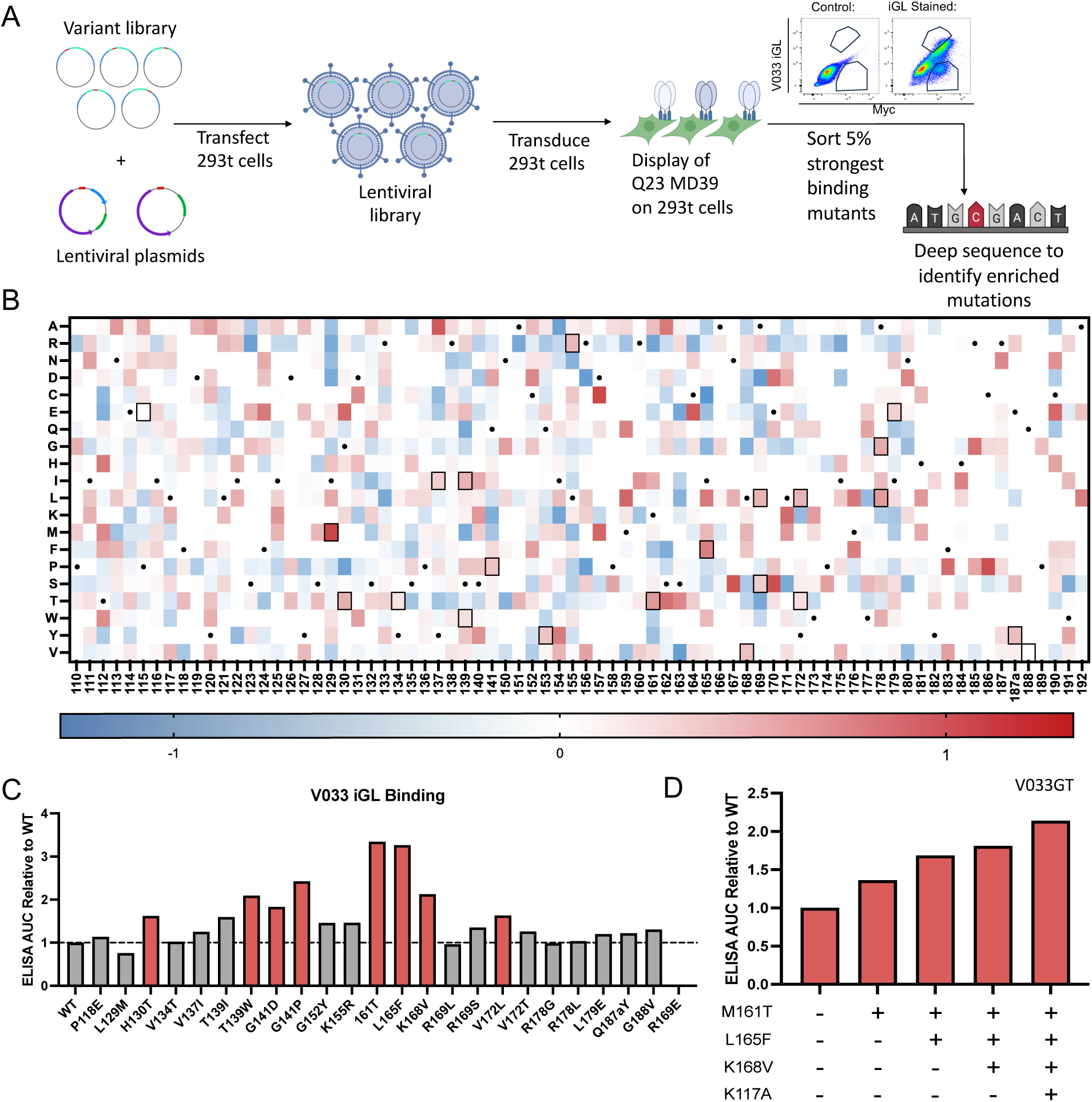
A single-round mammalian display approach for the identification of V033 iGL affinity enhancing mutations. **A.** Overview of our mammalian display approach. An immunogen variant library is packaged into lentiviral vectors, following which 293T cells are transduced with the library. Cells displaying the immunogen variant library are then stained with an antibody of interest, sorted, and deep sequenced to identify enriched mutations. **B.** Positive and negative enrichments following a single round of sorting against the V033 iGL. Black dots indicate the WT residue at each position. Boxes indicate mutations evaluated in (**C**). **C.** ELISA area under the curve binding values of the V033 iGL to Q23.MD39 site-directed mutants incorporating mammalian-display identified mutations. AUC values are normalized to WT. **D.** ELISA area under the curve binding values of the V033 iGL to Q23.MD39 mutants sequentially incorporating mammalian-display identified mutations. AUC values are normalized to WT.

As proof-of-principle, we sorted our Q23 library following staining with the V033 iGL. Sequencing of sorted cells led to the identification of mutations spanning the length of the library that were either enriched or selected against, providing an array of potential affinity enhancing mutations (Fig. 5B). From these enrichments, we filtered positions with low numbers of counts among either the selected or unselected cells, enrichments that arose from low numbers of a mutation appearing in the unsorted library, mutations that occurred at positions where the WT amino acid was also enriched, and mutations that altered or introduced cysteine or proline residues.

A subset of 24 especially enriched mutations that passed these criteria were synthesized as soluble Q23.MD39 trimer and then expressed, purified and tested for enhanced binding against the V033 iGL. We observed that the majority of the mutations conferred increased binding to the antibody by ELISA ranging from low (<1.5 fold) to high (>3 fold) increase in binding (Fig. 5C). We next asked whether these mutations had an additive enhancement of binding affinity when combined. We selected high binding mutations, M161T, L165F, and K168V and progressively incorporated them into Q23.MD39. ELISA binding to the V033 iGL showed increases in affinity as each mutation was incorporated (Fig. 5D). Subsequent addition of a K117A, identified through sorts against other axe-like antibodies, further increased the affinity of the V033 iGL, ultimately resulting in a construct named Q23.V033GT. We assessed binding against V033 iGL by SPR and found that the unmodified Q23.MD39 did not have measurable affinity (up to 1uM), however, Q23.V033GT bound at a K_D_ of 100nM (Fig. S5).

### A germline targeting SHIV elicits axe-like V2 apex antibodies

We next asked whether our Q23.V033GT iGL-targeted immunogen would be able to elicit structurally homologous antibodies in rhesus macaques. To test this, we introduced our germline targeting mutations into a replicating SHIV (70, 71) for the following reasons: (i) the high and sustained antigenic loads during viral infection maximize the probability of engaging bnAb precursor B cells, similar to extended dosing or using nucleic acid delivery; (ii) SHIVs coevolve with the bnAb lineage, selecting for affinity maturation and the acquisition of neutralization breadth, thereby allowing us to verify that primed B cells are genuine bnAb precursors; (iii) the pattern of Env escape can aid in the design of boosting immunogens, since escape variants select for affinity maturation and neutralization breadth; and (iv) sequencing of Env escape serves as a sensitive indicator of nAb-targeted epitopes, allowing identification of both on- and off-target nAb lineages (23, 72).

To construct this germline targeting SHIV, we first sought to incorporate our mammalian display-identified mutations into the WT SHIV-Q23 (70). We found that the V033-targeting mutations in Q23.V033GT could be incorporated into the virus without impacting viral infectivity (Fig. S6A). A SHIV bearing all 4 of these mutations, SHIV-Q23.V033GT maintained infectivity and p27 antigen titers similar to WT (Fig. S6A).

Strikingly, neutralization assays showed an over 50-fold increase in neutralization potency by the V033 iGL against the resulting SHIV-Q23.V033GT, verifying that the germline targeting mutations retained their effect in the context of an infectious virion (Fig. 6A). Antigenic profiling of the germline targeting SHIV demonstrated that the Q23.V033GT Env retained a closed prefusion state, showing no sensitivity to a panel of CD4i and V2p antibodies consisting of CH58, CAP228-3D, 447-52D, 17b, A32, 697-D, and 1393. However, the SHIV showed a modest increase in neutralization sensitivity to the antibody 3074, suggesting a more exposed V3 loop (Fig. 6A).

**Figure 6.**
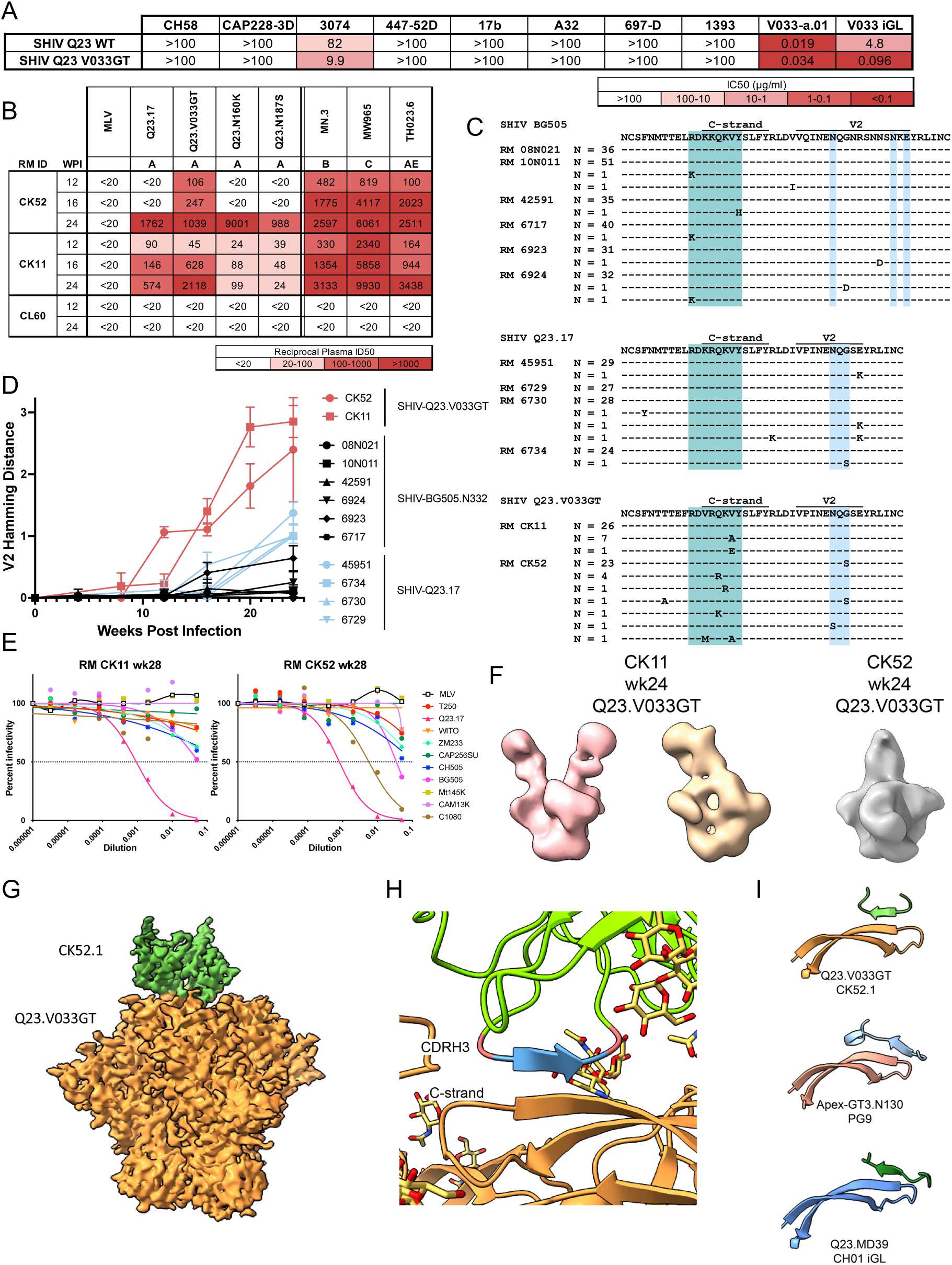
Infection with a germline-targeting SHIV elicits axe-like antibodies to the V2 apex. **A.** Neutralization IC50 values of SHIV-Q23.17 and SHIV-V033GT against a panel of non-neutralizing antibodies, the mature V033-a.01 bnAb, and the V033 iGL. **B.** Longitudinal plasma ID50 titers against MLV, autologous mutants, and tier-1A viruses. MLV, Murine Leukemia Virus. **C.** Single genome sequencing of circulating plasma viruses at week 12 post-infection in rhesus macaques infected with SHIV-BG505, SHIV-Q23.17, or SHIV Q23.V033GT. C-strand residues that are the primary site of Env escape from V2 apex bnAbs are highlighted in teal. V2b residues that are potential sites of glycan additions are highlighted in blue. N = number of viruses with a particular sequences. **D.** Longitudinal Hamming distances of V2 residues C131-C198 in the RMs from **C**. Error bars indicate a 95% confidence interval. **E.** Neutralization ID50 curves of week 28 plasma from macaques CK11 and CK52 against a panel of heterologous viruses. **F.** 3D reconstruction of selected NS-EM 3D classes that displayed V2 apex binding antibodies for macaques CK11 and CK52 at week 24. **G.** Cryo-EMPEM reconstructed density map of a V2 apex antibody (green) binding to Q23.V033GT (orange). **H.** Close-up view of the epitope recognized by the CK52 antibody CDRH3. Turn residues are colored red, while residues with beta strand backbone geometry are colored blue. **I.** Comparison of the CK52.1 CDRH3 with the mature PG9 bnAb (PDB: 7T77) and the CH01 iGL.

Three rhesus macaques were infected with SHIV-Q23.V033GT, as previously described (23, 70). All rhesus macaques were productively infected, with peak viral loads between 10^7^-10^8^ vRNA copies/ml of plasma 2 weeks post infection (Fig. S6B). Viral loads for one rhesus macaque, CL60, were especially high during the first 12 weeks of infection (>5x10^6^ vRNA copies/ml). This animal failed to develop any detectable antibody response to SHIV infection and succumbed to rapid clinical progression of AIDS (73) (Fig. S6B). The other two macaques exhibited stable setpoint viral loads between 5×10^4^–5×10^5^ vRNA copies/ml (Fig. S6B). Each mounted autologous Q23.V033GT and tier 1A nAb responses by 12 weeks post infection (Fig. 6B). Interestingly, the entire nAb response of rhesus macaque CK52 at week 12 was immunofocused to the V2 apex, as the response was dependent on the introduced germline targeting mutations (Fig. 6B). In both CK11 and CK52, we measured neutralization of V2 apex site-directed mutants (N160K and N187S) and observed striking changes in neutralization, thus confirming the presence of V2 apex targeted nAb responses (Fig. 6B). Single-genome sequencing of the circulating plasma virus quasispecies in CK52 and CK11 showed strong selection at the V2 apex at week 12 post-infection, with the addition of glycans to the V2b hypervariable loop and selection for neutralization escape at the V2 apex C-strand (Fig. 6C, Fig. S6D-E).

To understand how efficiently these C-strand targeted nAbs were being primed and affinity matured by SHIV-Q23.V033GT compared to SHIVs bearing wild-type Envs, we compared the kinetics of V2 apex targeted nAb elicitation in CK52 and CK11 with 6 macaques infected with WT SHIV-BG505.N332 and 4 macaques infected with WT SHIV-Q23.17 over the first 24 weeks of infection (Fig. 6C-D). We used kinetics of epitope escape as a measure of V2 apex-targeted nAb elicitation, since virus escape has been shown to be a sensitive and specific indicator of nAb responses (24, 74–76). To quantify escape, we calculated Hamming distance sequence similarity measurements in the V2 apex epitope relative to the starting SHIV sequence. At week 12 post infection, SHIV-Q23.V033GT infected macaques exhibited significantly more selection at the V2 apex than the SHIV-BG505.N332 (p = 0.0102) or SHIV-Q23.17 (p = 0.0170) infected macaques (Fig. 6C-D, p values determined by one-way ANOVA followed by Tukey’s post-hoc test). The acceleration of selective pressure on the V2 apex in SHIV-Q23.V033GT infected macaques was even more apparent at week 24 (Fig. 6D, p < 0.0001 against both BG505.N332 and Q23.17 by one-way ANOVA followed by Tukey’s post-hoc test). These data suggest that infection with SHIV-Q23.V033GT resulted in significantly faster and stronger selection at the V2 apex as compared to infection with primary HIV Envs, highlighting the ability of this Env to immunofocus early responses to the V2 apex. These findings suggest that a germline targeted SHIV with enhanced affinity for V2 apex precursors can consistently elicit V2 apex C-strand targeted nAbs early in infection.

To determine if the macaques with V2 apex responses also generated neutralization breath, we measured neutralization activity against a panel of nine heterologous tier-2 viruses. Limited heterologous neutralization was detectable in both macaques CK11 and CK52 at 28 weeks post-infection. Both plasma samples neutralized BG505 at titers of 1:20 to 1:30, and CK52 plasma also neutralized C1080 at a titer of 1:200 (Fig. 6E). Finally, we employed a powerful imaging technique, Epitope Mapping by Polyclonal Electron Microscopy (EMPEM), using both negative stain (nsEMPEM) (77) and cryo-EM (cryoEMPEM) (78) to confirm the presence of V2 apex directed antibodies in SHIV immune sera. The nsEMPEM data revealed strongly epitope focused responses, as we observed only V2 apex and gp41 binding antibodies at week 24 in macaque CK11 and only V2 apex antibodies in macaque CK52 (Fig. 6F, Fig. S7). Further, we observe multiple V2 apex targeting antibodies with an off-center angle of approach characteristic of axe-like bnAbs. By fitting high resolution Env-axe antibody complexes into the CK52 nsEMPEM density, we observed significant overlap with that of the V033, 41328 and CH01 iGL Fabs, indicating the presence of a structurally homologous antibody (Fig. S7C). Next, we performed cryo-EMPEM on the CK52 sample at week 24 and obtained a 4.2Å resolution structure (Fig. 6G). Consistent with nsEMPEM results, the cryo-EMPEM confirmed V2 apex engagement targeting the C strand region similar to the approach of axe-like bnAbs (Fig. 6H). Further refinement of antibody CK52.1 revealed turn-beta-turn density likely contributed by the CDRH3 interacting with the C-Strand as expected for axe-like engagement of Env V2 apex (Fig. 6H, 6I). Together, the escape mutations, neutralization sensitivity and structural data demonstrate that SHIV-Q23.V033GT re-elicited axe-like antibodies to the Env V2 apex early in infection.

### Construction of an axe-targeting immunogen

We next sought to simultaneously enhance the affinity of Q23.MD39 for multiple known axe-like iGL antibodies. We reasoned that an immunogen with increased affinity for multiple immunogenetically distinct iGLs of this class would be generally more efficient in binding naïve germline B cell precursors harboring this CDRH3 topology. To do this, we took advantage of the large number of enrichments identified by our mammalian display approach. By generating matrices of enrichments following sorting of our NNK library against the CH01, PG9, and V033 iGLs and looking for enrichments in common, we were able to identify a set of mutations that simultaneously increased affinity for the V033, PG9, and CH01 iGLs, as well as to the 41328 iGL which was not included in our mammalian display staining (Fig. 7A-B). The mutations in this trimer, Q23.RH-GT (for Rhesus-Human Germline Targeting), were all distal to the C-strand, the primary point of contact at the epitope-paratope interface (Fig. S8), and instead may have enhanced affinity through second shell interactions or indirect restructuring of the Env apex, making it more amenable to engagement by axe-like antibodies.

**Figure 7.**
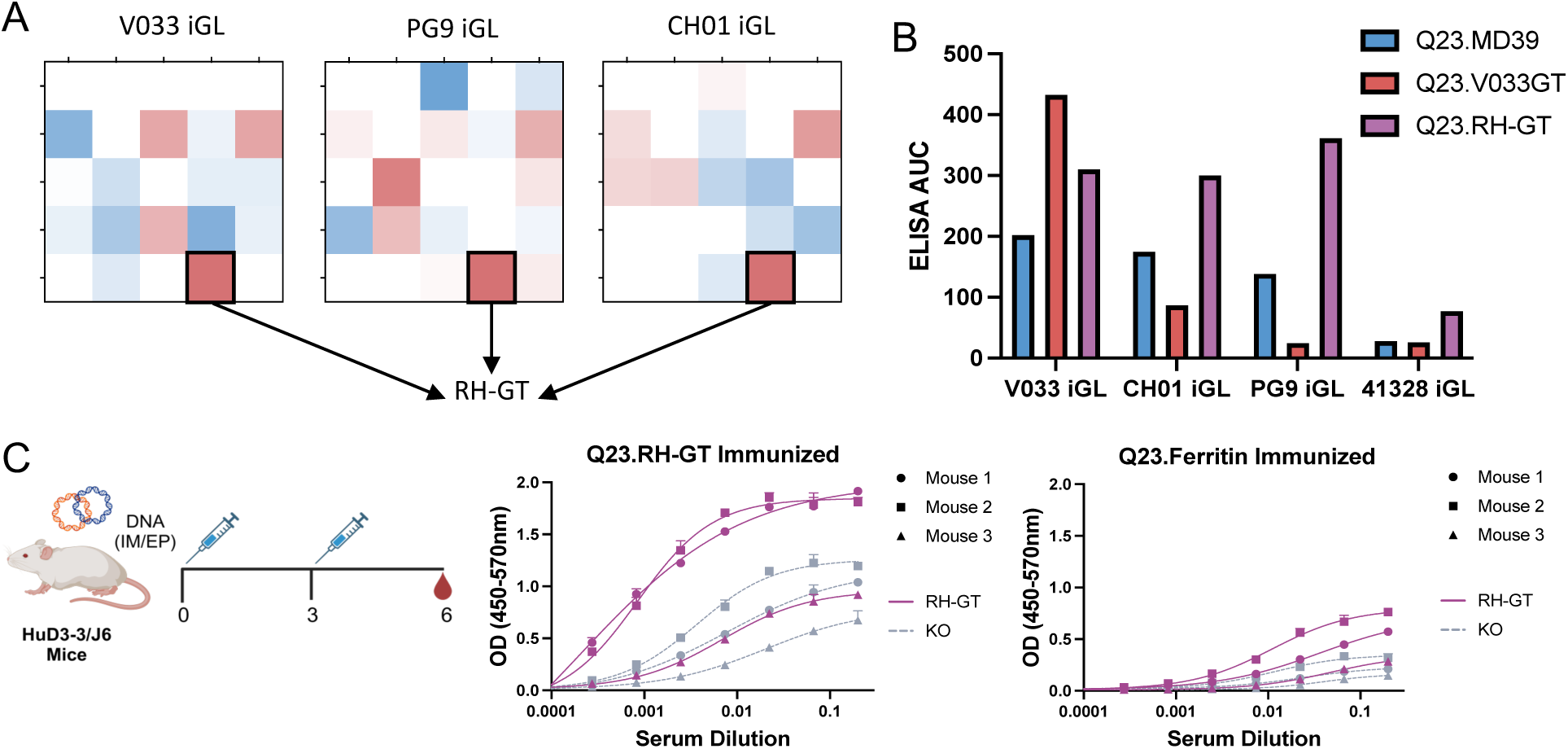
Generation of a multi-axe targeting immunogen. **A.** Overview of our approach for immunogen design. Mutations enriched in multiple mammalian display mutagenesis sorts against a several antibodies belonging to a single structural iGL class are identified and incorporated into an immunogen. **B.** ELISA area under the curve binding of Q23.MD39, Q23.V033GT, and Q23.RH-GT against axe-like V2 apex bnAb precursors. **C.** (Left) HuD3-3/J6 mice were primed and boosted with 25µg of DNA-encoded Q23.RH-GT adjuvanted with 0.5µg of IL-12 plasmid. (Right) ELISA binding of serum from mice immunized with Q23.RH-GT or WT Q23.Ferritin to Q23.RH-GT or a Q23 epitope knockout. Shapes indicate individual mice. Solid purple lines indicate binding to Q23.RH-GT. Dashed grey lines indicate binding to Q23.KO.

To confirm that Q23.RH-GT still maintained the fidelity of its V2 apex epitope as well as an ability to engage with axe-like antibodies, Cryo-EMPEM was used to investigate the ability of this Env to bind Fabs from the polyclonal plasma of CK52. This yielded a 3.78 Å Cryo-EM density map that exhibited a V2 apex Fab density that aligned remarkably with the CK52 Fab density observed in complex with Q23.V033GT. The V033-a CDRH3 resulted in a well-fit peptide segment in the CK52/Q23.RH-GT Cryo-EMPEM density map (Fig S9). Together, these results show that Q23.RH-GT exhibited the ability to engage with a heterologous axe-like precursor and maintained the structural architecture of the V2 apex epitope after iterative rounds of mammalian display.

To test the ability of this immunogen to elicit axe-like bnAb precursor lineages, we immunized hD3-3/JH6 mice (79). This model has the mouse DQ52 and JH1-4 segments replaced by human IGHD3-3 and IGHJ6 segments, respectively, allowing the B cells to incorporate these genes during VDJ recombination and produce diverse CDRH3s of lengths longer than WT mice and comparable to human V2 apex bnAbs that incorporate these same genes (79). The hD3-3/JH6 mice (n=3) were immunized with 25µg of DNA encoding an RH-GT trimer along with IL-12 adjuvant and boosted 3 weeks post prime (Fig. 7C). All mice showed strong binding to the Q23.RH-GT trimer but diminished binding to a V2 apex C-strand epitope knockout trimer (Q23.MD39 R169E/K71E), indicating the presence of robust C-strand directed responses (Fig. 7C). Control mice immunized with Q23.ferritin immunogen containing a WT V2 apex epitope. These mice failed to induce the same level of epitope-specific response. These data indicate that the Q23.RH-GT immunogen was successfully able to induce V2 apex responses in a stringent rearranging mouse model.

## Discussion

While the precise structures of antibody CDRs are crucial for understanding their specific interactions with antigens, our knowledge of the structural landscape of B cell receptors within the human antibody repertoire remains limited. A central hypothesis of the present study is that AI and structural informatics can be used to augment germline targeting vaccine design strategies through the identification of broader pools of desired antibody precursors. As proof-of-principle of this approach, we apply it to the HIV Env V2 apex epitope by determining the frequency of antibodies in the human repertoire with a specific CDRH3 structure, designing immunogens capable of binding to antibodies harboring these CDRH3 structures, eliciting neutralizing antibodies with this target CDRH3 topology in non-human primates through infection with a germline targeting SHIV, and eliciting epitope-specific responses in humanized mouse models. Our structurally focused approach to precursor targeting is predicated on three key observations. First, our structure of the CH01 iGL retains the distinct CDRH3 topology of mature lineage members, as was previously seen with the V033-a.UCA (24), PCT64 LMCA (25), and PG9 iGL (25) precursors. This suggests V2 apex bnAb precursors often acquire their distinct CDRH3 conformations during VDJ recombination, and not by affinity maturation, and therefore bnAb precursors may be identified through these distinct CDRH3 conformations. Second is the isolation of rhesus V2 apex-targeting antibodies that utilize the rhesus-specific D3-15*01 gene (32–35) but nonetheless recapitulate the axe- and needle-like shapes found in C-strand–targeting human V2 apex bnAbs. This structural homology, combined with the immunogenetic diversity of human V2 apex bnAbs, underscore the ability of immunogenetically distinct CDRH3s to adopt convergent solutions to epitope recognition. Third, we show that structure prediction methods, such as AF2, are capable of predicting V2 apex bnAb CDRH3 structures, as illustrated by its ability to accurately predict the CDRH3 topologies of rhesus V2 apex bnAbs absent from its training dataset (Fig. S2). This indicates that AI-based structure prediction can be used to identify B cells with particular CDRH3 topologies in an automated fashion, as demonstrated by our screens for B cells bearing a structurally conserved ‘turn-beta-turn’ CDRH3 topology.

A surprising finding of the precursor analyses in the present study is the identification of a significant pool of B cells in the human repertoire that resemble rhesus V2 apex bnAb precursors. Several studies have identified the rhesus IGHD3-15*01 gene, which lacks a human homolog, as nearly universally incorporated into rhesus V2 apex bnAbs (32–35, 72). Structural analysis of a key “EDDYG” motif within antibody CDRH3s from this D gene showed that it played diverse roles in epitope engagement and therefore was strongly selected for its unique biochemical features (32), rather than for its ability to reproducibly engage specific epitope residues as seen previously for other highly selected immunogenetic elements (80, 81). This, however, raises the question as to whether the presence of this favorable rhesus-specific D gene makes rhesus V2 apex bnAb precursors inauthentic representations of B cell precursors in humans. This concern is allayed by our repertoire analysis in which we identified B cells with the key “EDDYG” motif and suitable CDRH3 lengths at frequencies as high as 1 in 100,000 B cells in every donor. It will thus be intriguing to see the results of upcoming human vaccine trials of germline-targeting V2 apex immunogens, which will enable a direct comparison of V2 apex-targeted responses between humans and rhesus macaques.

The identification of B cells bearing the “EDDYG” motif in the human repertoire supports the use of rhesus V2 apex bnAb precursors for human vaccine engineering, exemplified by our Q23.RH-GT immunogen which exhibited increased affinity for both rhesus and human precursors. Notably, several other investigators have used rhesus bnAb precursors to design V2 apex immunogens and have obtained promising immunogenicity results with respect to the elicitation of V2 apex bnAbs. These include the multivalent display of Env on liposomes (34) and trimers engineered through structure and evolution guided designs that exhibit enhanced binding affinity for multiple rhesus and human V2 apex bnAb precursors (35). Ma and colleagues also used evolution-guided mammalian display mutagenesis to identify an engineered Env trimer, ApexGT6, that successfully induced V2 apex bnAb-like precursors in rhesus macaques (33). Altogether, these results support the V2 apex as a highly promising target for germline-targeting strategies and the Q23.17 Env as a strong candidate platform for the design of such immunogens.

Interestingly, many of these recent V2 apex-targeting immunogens also used the Q23.17 Env as a vaccine platform. Previously, we showed that the wildtype Q23.17 Env, compared with 15 other primary HIV-1 Envs, was the most efficient in inducing V2 apex bnAbs in macaques when expressed as replicating SHIVs (24). We further observed that a single prime and boost with this Env expressed as soluble protein trimers could elicit broadly neutralizing antibodies in a rhesus V2 apex bnAb UCA knock-in mouse model (82). Several features of our structure of Q23.MD39 in complex with the CH01 iGL help explain these observations. First, the C-strand of Q23.17 is enriched in basic residues, creating a positively charged surface that complements the negatively charged CDRH3s of most V2 apex bnAb precursors (24). Second, Env Q23.17 lacks glycans at residue 130 and in the V2b loop, both of which can occlude access to the V2 apex by germline antibodies. Third, the length of the V2b loop is short enough to minimize clashes with the approaching Fab, enabling binding by relatively low affinity antibodies. Our results here show that germline-targeting mutations can be engineered into Env Q23.17 to elicit V2 apex-targeted responses in humanized mouse models, and these mutations can elicit in rhesus macaques heterologous tier-2 neutralizing antibodies that harbor the turn-beta-turn axe-like CDRH3 identified in our structural analysis of the B cell repertoire. With the advent of more accurate protein structure prediction tools, we expect our structure-guided immunoinformatic approach to provide an increasingly powerful method for precursor analyses. Furthermore, we anticipate that this approach will open the door for new studies on other bnAb epitopes on HIV Env as well as other glycoproteins with rich sets of antibody structures such as SARS-CoV-2, influenza, and RSV.

## Acknowledgements

We would like to thank Drs. Frederick Alt and Ming Tian for provision of the human D3-3/J6 mouse model. In addition, we would like to thank the Wistar Animal, Molecular Screening & Protein Expression, and Flow Cytometry Facilities for experimental support. Funding was provided under NIH grants U19 AI166916 (D.B.W.) and R01 AI165080 (G.M.S.). R.H is supported by a Training Grant in HIV Pathogenesis (T32-AI007632). S.O.S. is supported by a Training Grant in Structural Biology and Molecular Biophysics (T32-GM132039). We thank Thomas Klose at the Purdue University Cryo-EM facility, Ashok Nayak at Thomas Jefferson University, and Sudheer Molugu and Prerana Gogoi at Beckman Center for Cryo-EM at the University of Pennsylvania for their assistance with data collection.

## Materials and Methods

All materials and methods are described in detail in the *SI Appendix*.

### Data, Materials, and Software Availability

The atomic models of the 35O22/Q23.MD39 and CH01 iGL/35O22/Q23.MD39 complexes generated in this study are available at the Protein Data Bank (PDB, https://www.rcsb.org) under the PDB accession codes 9DIM, 9DHW, and 9Z9L. The corresponding cryo-EM reconstruction is available at the Electron Microscopy Data Bank (EMDB, https://www.ebi.ac.uk/emdb/) under the EMDB access codes 46914, 46884, and 73949, respectively. Cryo-EMPEM reconstructions are available at the EMDB access codes 73950 and 73961. Longitudinal Env SGS gp140 sequences are deposited at Genbank under accession numbers MN467402–MN468119, PX382176–PX382323, MN468931–MN469066, PX301680–PX302024, and PX384478–PX384772.

