## Supplemental Information for "Deep Mining of the Human Antibody Repertoire Identifies Frequent and Genetically Diverse CDRH3 Topologies Targetable by Vaccination"

### Supporting Information Text

### Extended Methods

#### Inferred germline precursor (iGL) inference

iGLs for rhesus bnAb lineages 41328-a and V033-a were constructed by reverting V, D and J gene templated amino acids to their germline gene sequences. Non-templated CDRH3 residues were left unchanged from the mature bnAb sequence (1). For human bnAb lineages CH01 and PG9, previously reported iGLs were used (originally referred to as CH01 UA3 and PG9 UA respectively) (2).

#### Repertoire frequency analysis

Our antibody searches used the heavy chain only sequences from healthy, HIV-unexposed human donors. This resulted in over 1 billion sequences. We calculated the B cell frequencies within each donor for each search by taking the number of sequence hits and dividing that by the number of sequences the donor had in the OAS database. We then removed redundant sequences within each donor. For each donor, we first grouped the donor's sequences into groups based on CDRH3 length. Then, we compared CDRH3 sequences in a pairwise fashion for each length group. If the number of different residues between two CDRH3s was less than 15% of the CDRH3 length (greater than 85% similarity within CDRH3) then the sequence was considered redundant and was removed. For a CDRH3 of length 20, this would be a difference of 3 or fewer amino acids and for length 40, a difference of 6 or fewer amino acids. Our exact searches and example scripts are deposited in the GitHub repository: https://github.com/ssolieva/q23-manuscript.git

#### AlphaFold2 modeling

We generated 1 seed per model using ColabFold (3) (AlphaFold2-multimer (4) with custom templates for our structure predictions. We created custom templates by removing the CDRH3 loop from the CH01, V033, and PG9 structures. The custom templates were used without a multiple sequence alignment and the CDRH3 region was untemplated. Each antibody was modeled in about 6 minutes. We paired our non-redundant variable heavy chain sequences from the OAS searches with the mature variable light chain from the antibody the search was based on (CH01, PG9, V033).

#### Structural axe definition

We created a structural definition of an axe using the four available axe antibodies. We defined an axe as having the following pattern: at least two turn residues, followed by at least two beta residues at least 20 A from the start of the CDRH3, followed by at least two turn residues. We defined a turn residue as having any of the following: (IDSSP-defined hydrogen-bonded turn or bend, (ii) hydrogen bond between residues *i* and *i*+*3* or between residues *i* and *i*+*4, or* (iii) Phi/Psi angle-defined Type I, I', II, or III turn (see table below). We defined a beta residue as having any of the following: (i) DSSP-defined beta sheet (residue in isolated beta-bridge or extended strand), or (ii) Phi/Psi angle-defined beta sheet (phi < -50 and psi > 95 or phi < -50 and psi < -170). The beta residue was defined as a 'far beta' if the residue's alpha carbon was at least 20 A away from the alpha carbon of the first residue in the CDRH3.

| Phi/Psi angle definitions for turn types I, I', II, and III. | | |
|---|---|---|
| Turn Type | Residue i+1 | Residue i+2 |
| Type I | Phi: -100 to -25<br>Psi: -75 to 25 | Phi: -150 to -50<br>Psi: -50 to 50 |
| Type I' | Phi: 25 to 100<br>Psi: 0 to 75 | Phi: 50 to 150<br>Psi: -50 to 60 |
| Type II | Phi: -100 to -25<br>Psi: 75 to 180 | Phi: 50 to 125<br>Psi: -50 to 50 |
| Type III | Phi: -100 to -50<br>Psi: -60 to 10 | Phi: -150 to -60<br>Psi: 75 to 160 |

To validate our structural axe definition, we applied it to 610 human antibody structures from the SAbDab (5). The specific structures we used and their classifications using our definition can be found in our GitHub repository: https://github.com/ssolieva/q23-manuscript.git

**MD simulations**

Molecular dynamics simulations were seeded from AlphaFold2-generated, cryo-em, or x-ray crystallography structures using the GROMACS software (6) (version 2023.2) and the CHARMM36m force field (7). The structures consisted of a variable heavy chain and the mature variable light chain for the respective bnAb (CH01, V033). The antibodies were each solvated using TIP3P waters (8) and 0.1 uM NaCl in a dodecahedron box. Energy minimization was performed using steepest descent minimization, followed by NVT equilibration for 200 ps, NPT equilibration for 1 ns, and 10 independent 250 ns production runs per antibody, with snapshots saved at 20 ps intervals. We calculated the frequency of axe-like structures throughout each trajectory by applying our axe structural definition at 200 ps intervals. This analysis was conducted using the MDTraj software (9). All input files, starting structures, and analysis files are deposited in the following GitHub repository: https://github.com/ssolieva/q23-manuscript.git

***In silico* saturated mutagenesis**

In silico saturated mutagenesis was conducted using AlphaFold2 and mutating each residue on the CH01 CDRH3 to every possible amino acid. Each prediction had 5 models and 10 seeds per model, resulting in 50 structures per mutation. The RMSD values were computed between the CDRH3 regions of the CH01 AlphaFold2 structure and the AlphaFold2 predictions of the mutations. The analysis files are deposited in the following GitHub repository: https://github.com/ssolieva/q23-manuscript.git

**DNA design and plasmid synthesis**

The amino acid sequence for Q23.17 was obtained from a previously published sequence (genbank accession number AF004885.1). Stabilized trimer constructs were developed by adding published stabilization mutations into the Q23.17, and all constructs included the T533A “repair” mutation. Constructs were then codon optimzed, and an optimized IgE leader sequence was added to the N terminus of the protein to provide efficient processing and secretion. INO-9012 IL-12 DNA (pIL-12), the cytokine adjuvant is a single plasmid, plasmid IL-12 was admixed with pQ23.MD39 or pQ23.Ferritin at appropriate amounts prior to IM administration. All plasmids were synthesized and cloned (GenScript) into our modified pVax1 backbone (Inovio Pharmaceuticals).

**Antibody expression and purification**

Expi293F cells (ThermoFisher) were maintained in Expi293 expression medium (ThermoFisher). All cell lines were mycoplasma negative and tested on a regular basis. All proteins were produced by Expifectamine transfection of Expi293F cells following the manufacturer’s protocol. Transfection enhancers were added 18 h after transfection and supernatants were harvested 6 days later. Antibodies were purified using Protein A agarose according to the manufacturer’s protocol to purify the IgG. Purity was confirmed with Coomassie staining of SDS-page gels and concentration was determined using a nanodrop.

**Pseudovirus production and purification**

Pseudotyped viruses were produced using HEK 293 T cells transfected with 4 ug of a plasmid expressing the Env of interest and 8 ug of a plasmid expressing the HIV backbone Δ Env (pSG3ΔEnv − NIH AIDS Reagents) using GeneJammer (Aglient). Forty-eight hours after transfection, cell supernatant was harvested, filtered through a 45 μm filter, aliquoted, and stored at −80 °C.

**Trimer production and purification**

Env-based trimers were expressed in Expi293F cells maintained in Expi293 expression medium (ThermoFisher). Trimers were produced by Expifectamine transfection (Gibco, A14524) of Expi293F cells following the manufacturer's protocol. The trimer-containing supernatants were obtained by centrifuging (4000 × g, 25 mins) and filtering (0.2 um Nalgene Rapid-Flow Filter) the 293F cultures, following which trimers were purified from supernatants by lectin purification using lectin beads (Vector Laboratories) and lectin elution buffer (1M Methyl alpha-D-mannopyranoside). The trimers were then purified over a size-exclusion chromatography column (GE S200 Increase) in PBS. The molecular weight and homogeneity of the trimers were confirmed by protein conjugated analysis from ASTRA with data collected from a size-exclusion chromatography-multi-angle light scattering (SEC-MALS) experiment run in PBS using a GE S6 Increase column followed by DAWN HELEOS II and Optilab T-rEX detectors. The trimers were aliquoted at 1 mg/ml and flash frozen in thin-walled PCR tubes prior to use.

**Antigen Conformation Tracing in vivo by ELISA (ACTIVE)**

For Antigen Conformation Tracing *in vivo* by ELISA, BALB/c mice were administered with 100 µg DNA plasmid co-formulated with 12 U hyaluronidase in the tibialis anterior (TA) muscles of the mice as described previously (10). At predetermined time points TA muscles were harvested and homogenized in T-PER extraction buffer (Thermo Fisher Scientific) containing protease inhibitor (Roche). Muscle homogenates were subsequently concentrated using a 3kDa Amicon Ultra 0.5 mL centrifugation kit (Millipore Sigma). Concentrations of total proteins were estimated using BCA assay kit (Thermo Fisher Scientific). In order to determine correct folding and binding of *in vivo* expressed antigens, 96 well ELISA plates (Corning, 3690) were coated with 4 ug/ml of recombinant PGT128 Fab fragments in PBS and incubated overnight at 4°C. After washing, plates were blocked with 5% skimmed milk in PBS containing 1% newborn calf serum (NBS) and 0.2% Tween for 1 hour at RT. Total protein concentrations were normalized and added to the plate followed by serial dilutions. Recombinant BG505.MD39, Q23.MD39, Q23.Ferritin and gp120-foldon were added as positive control standards. Plates were incubated at RT for 2 h, then followed by washing, antibodies of interest were added at 10 ug/ml except for PGT145 which was added at 50 ug/ml, for 1 h at 37 °C. Plates were further washed and incubated with Goat Anti-Human IgG Fc Fragment conjugated with HRP (Bethyl Laboratories Inc) for 1hour at RT. Plates were developed for 5 min with 1-step ultra TMB (Thermo Fisher Scientific) and stopped with 1 N $H_2SO_4$. Absorbance at an optical density (OD) of 450 nm and 570 nm was measured using Synergy2 plate reader (BioTek Instrument). The background 570 nm OD was subtracted from the 450 nm reading. The data was analyzed and fitted using Graph Pad Prism 10.2.

**Antibody Digestion for Complexation**

Monoclonal and polyclonal antibodies were digested into antigen-binding fragments (Fabs) by adding antibodies into 6-12 mL of digestion buffer (100 mM sodium acetate 10 mM L-cysteine 0.3 mM EDTA pH 5.6 or 7) followed with the addition of papain (2%, w/w), pre-incubated for 15 mins in digestion buffer. Digestion reactions were incubated overnight and then quenched using 3 mM iodoacetamide. Protein A resin was added into the quenched digestion mixture and incubated for 15 mins on ice. Fab was purified from the Protein A/digested antibody mixture through a gravity column. Flow through fraction was collected and buffer exchanged into 1X PBS using Amicon-Ultra concentrator with molecular weight cutoff of 10 kDa. Protein A was resin washed with at least 10 CV of 1X PBS and eluted using protein A elution buffer (20 mM citrate, 100 mM NaCl, pH 3). Flow through, wash, and elution fractions were analyzed by SDS-PAGE. CH01iGL Fabs were unable to be purified using Protein A resin due to binding of Fabs to Protein A. Instead, digestion mixture mixed with 3mM iodoacetamide was buffer exchanged into 1X PBS. CH01 iGL Fabs/Fc/IgG mixture was separated using Superdex 200 increase 10/300 size exclusion column using 1X PBS as the running buffer to obtain the CH01iGL Fabs/Fc mixture. Fractions containing CH01iGL Fabs/Fc were concentrated using Amicon-Ultra concentrator with a molecular weight cutoff 10kDa. All Fabs were stored at 4°C.

**Negative stain electron microscopy sample preparation and data collection**

SEC purified MD39 trimers were further dialyzed into Tris-buffered saline (TBS). A total of 4 µL of purified proteins (0.005 mg/mL) was applied onto glow discharged carbon-coated Cu400 EM grids.

The grids were then stained with 4 µL of 2% (w/v) uranyl formate, blotted, and stained again with 4 µL of the stain followed by a final blot. Image collection was performed on a FEI Tecnai T12 microscope equipped with Oneview Gatan camera at 62,750x camera magnification resulting in pixel size of 2.356 Å.

For nsEMPEM, Q23.V033GT (50 µg) was mixed with 500 µg of polyclonal fabs and incubated at 4 degrees overnight. The resulting complex mixture was purified using Superose 6 increase 10/300 size exclusion column and complex fractions were collected. The collected fractions were diluted immediately and applied onto glow discharged carbon coated 300 mesh Cu grids (Electron Microscopy Sciences (EMS) CFTH300-Cu-50) to incubate for 2 mins before blotting. Complexes were stained using 2% uranyl formate. Micrographs were collected on a FEI Tecnai T12 microscope at 62,750x camera magnification resulting in pixel size of 2.356 Å/pixel.

**Negative stain electron microscopy data processing**
Micrographs were imported into RELION5.0 (11) and performed CTF estimation using CtfFind4. Particles were picked using Laplacian of Gaussian (LoG) picker and extracted for 2D classifications. Negative stain EMPEM data processing strategy from previous literature was implemented. (DOI: 10.1016/j.xpro.2023.102476) After the final round of 2D classification, there were 209,484 particles in the CK52 dataset and 59,730 particles in the CK11 dataset. These particles were then 3D classified with 50 classes for CK52 and 60 classes for CK11 (T=4). Good 3d classes were individually 3D refined using a 15 Å ligand-free Env density as a reference.

**Preparation of Q23.MD39 in complex with CH01 iGL and 35O22**
Q23.MD39/CH01 iGL/35O22 complex was prepared by mixing Fabs with a 1:40:40 molar ratio (Q23.MD39:CH01 iGL:35O22) in 1X PBS. Q23.MD39 complex was purified by Size-Exclusion Chromatography using the ENrich$^{TM}$ 650 column (BioRad) with 1X PBS as the running buffer and fractions containing the complex were collected and concentrated to 0.1 mg/mL. Purified protein complex was aliquoted for storage in -80°C until cryo-EM sample preparation.

**Cryo Electron Microscopy Sample Preparation, Data Collection, and Data Processing**
Q23.MD39/CH01 iGL/35O22 sample was diluted to 0.07 mg/mL and deposited on graphene-oxide (GO) coated Au-Flat grids (Protochips). Graphene oxide coating of Au-Flat grids was done in-house following a protocol from previous literature (12). 4 µL of sample was added to the GO-coated grid at 4°C under 100% humidity in a Mark IV Vitrobot (FEI), blotted with Whatman #1 filter paper, and then plunged immediately into liquid ethane. Dose-fractionated data for Q23.MD39/CH01 iGL/35O22 was collected in Counting mode at a magnification of 81,000x resulting in pixel size of 1.054 Å/pixel using aberration-free image shift (AFIS) protocol through the EPU software (ThermoFisher).

Data processing was performed employing a standard cryo-EM data processing workflow comprising motion correction, CTF estimation, reference-free LoG picking, 2D classification, manual inspection/selection of 2D class averages, and 3D refinement in RELION v3.1 (11). 3D classification was employed to remove junk particles as well as separating particles with or without CH01 iGL bound and with two and three 35O22-bound particles. Particles from two and three 35O22-bound Q23/CH01 iGL classes were individually 3D Refined and Bayesian Polished in RELION. Polished particles were pooled and transferred into CryoSPARC and focus 3D classified to further improve the CH01iGL density. The resulting 3D classes were individually refined, and the best class was manually inspected and used for another round of Non-Uniform Refinement to generate the final density maps.

**Cryo EMPEM Sample Preparation and Data Collection**
Q23.V033GT (100 µg) was mixed together with 1 mg of CK52 week24 polyclonal Fabs or 0.8 mg of CK11 week24 polyclonal Fabs and incubated at 4 degrees overnight. Polyclonal Fabs/Q23.V033GT complexes were purified using S6i 10/300 size exclusion chromatography in 1X PBS and fractions were collected, pooled, and concentrated using Amicon-Ultra concentrator with a molecular weight cutoff of 10 kDa. Complexes were diluted to 0.075 mg/mL and deposited

onto GO-coated UltrAuFoil R1.2/1.3 grids (Quantifoil). CK52week24/Q23.V033GT grids were loaded onto a Thermo Scientific Glacios (Thermo Fisher) equipped with a Falcon 4 detector. Dose-fractionated data was collected at 150,000x magnification resulting in a pixel size of 0.95 Å using AFIS protocol through EPU software (Thermo Fisher). Data was collected with a total dose of 50 $e^-/Å^2$ divided over 45 frames. CK11week24/Q23.V033GT grids were loaded on to Titan Krios Cryo Electron Microscope equipped with K3 Summit detector. Dose-fractionated data was collected at 105,000x magnification resulting in a pixel size of 0.43 Å in Counted Super Resolution mode using AFIS protocol through EPU software (Thermofisher). Data was collected with a total dose of 40.13 $e^-/Å^2$ divided over 37 frames.

**Cryo EMPEM Data Processing**

Micrographs were imported into cryoSPARC for motion correction and CTF estimation. Initial map reconstructed from blob picker was used for template picking. Particles from both picking strategies, with duplicated particles removed, were extracted (binned by 4), and 2D classified, and trimer particles were further filtered through two rounds of heterogenous refinement. Particles were re-extracted (binned by 2) and one round of Non-Uniform refinement was performed. Finally, particles were re-extracted with the original pixel size and were Non-Uniform refined using a 300 Å sphere mask. Particles used in the refinement were symmetry expanded through C3 symmetry. Symmetry expanded particles were 3D classified (K=10) using an 80 Å focus mask on the partial fab density and a 300 Å solvent mask. Particles from good classes were pooled then subjected to local refinement in CryoSPARC using a low-pass filtered map from one of the good 3D classes as a static mask. The resulting particles with further improved angular assignments were 3D-classified again using a 120 Å sphere focus mask around the Fab and the same solvent mask. A final round of local refinement was performed.

**Model Building**

A homology model of Q23.MD39 was generated from SWISS-MODEL (13) using a refined model of another high-resolution prefusion-closed Env trimer from the Pallesen Lab (unpublished). A homology model of CH01-iGL Fab was generated from SWISS-MODEL using CH03 Fab (PDB: 5ESV) (14). The crystal structure of 35O22 Fab (PDB: 4TOY) was used. Homology models were docked into the Q23.MD39/CH01 iGL/35O22 density map (UCSF ChimeraX) (15) and real-space refined using Coot (16). In-house developed protocols using Rosetta was employed for model refinement. N-linked glycans were added manually in Coot. Q23/35O22 atomic model was refined using the same protocol. Model-to-map fit was validated using EMRinger(17), glycan geometry using Privateer31(18), and model geometry using MolProbity (19).

**Serological trimer-binding ELISA**

Binding titers to trimer were determined by coating plates with 2ug/ml of recombinant PGT128 antibody overnight in PBS. After washing, plates were blocked with 5% skim milk in PBS with 1% newborn calf serum (NBS) and 0.2% Tween for 1 h at RT. Recombinant trimer was added at 4 ug/ml for 2 h at RT. Serum was serially diluted, added to plates, and incubated at 37 °C for 1 h. Antigen and species-specific IgG was then detected across absorbed secondary anti-mouse HRP antibody (Bethyl Laboratories Inc). Plates were developed for 5 min with 1-step ultra TMB (Thermo Fisher Scientific) and stopped with 1 N $H_2SO_4$. Absorbance at an optical density (OD) of 450 nm and 570 nm was measured on a Synergy2 plate reader (BioTek Instrument). The background 570 nm OD was subtracted from the 450 nm reading.

**Recombinant antibody binding ELISA**

Antibody affinity to Env trimers was determined by coating plates with 4 ug/ml PGT121 or PGT128 fab in 1×PBS for 3 h at RT. After washing with 1×PBS containing 0.05% Tween, plates were blocked overnight with 1×PBS containing 0.1% Tween and 5% skim milk. Plates were washed and trimer was incubated at 10 ug/ml for 1 h at RT. Plates were washed and recombinant antibody was serially diluted, added to plates, and incubated for 1 h at RT. After washing the plates, goat anti-human IgG Fc (Bethyl Laboratories Inc) at a dilution of 1:10,000 was incubated for 1 h at RT. Plates were washed and developed for 10 min with 1-step ultra TMB (ThermoFisher) and stopped with 1 N $H_2SO_4$. Absorbance at an optical density (OD) of 450 nm and 570 nm was measured on a Synergy2 plate reader (BioTek Instrument). The background 570 nm OD was subtracted from the 450 nm reading.

**DNA encoded immunogen immunization in WT (BALB/c) mice**

Female BALB/c mice of 6-8 weeks old were immunized with 5ug, 10ug and 25ug of plasmid DNA encoding Q23.MD39 trimer or 5ug and 10ug of Q23.Ferritin nanoparticle. The animals received these immunizations either with or without plasmid DNA encoding IL-12 as intramuscular (IM) injections into the tibia anterior (TA) muscles, followed by *in vivo* electroporation (EP) using the CELLECTRA ® -3P device (Inovio Pharmaceuticals). Mice were immunized at 0,3 and 6 weeks and sera were collected 2 weeks post each immunization through the submandibular vein for assessment of humoral immune responses.

**Neutralization Assays**

Pseudotyped viruses and SHIVs were titered on TZM-bl cells to determine IU/ml. All sera samples were heat-inactivated for 60 min at 56°C. 96 well plates were seeded with $10^4$ TZM-bl cells/well cultured in DMEM + 10% FBS one day prior to the assay. Serum was serially diluted in media supplemented with 10% normal human serum and incubated with pseudotyped virus and dextran (ThermoFisher) for 1 hour at 37°C, following which the virus/serum mixture was added to adherent TZM-bl cells. Forty-eight hours after incubation, media was removed and cells were lysed using PBS + 0.01% Triton-X (Promega). Luciferase luminescence was then measured using the Synergy2 plate reader (BioTek Instruments). Serum titer was determined for 50% virus neutralization (ID50).

**Mammalian display library design**

For cell surface display, Q23.MD39 was genetically fused to a PDGFR transmembrane domain via a G/S linker as previously described (20). This construct was synthesized in a pENTR backbone by Twist Biosciences. A scanning NNK library covering HXB2 residues 110-192 of the Q23.MD39 construct was produced using a BioXP 3250 instrument. NNK fragments were then pooled and assembled into the Q23.PDGFR.pENTR construct via Gibson assembly (NEB, E2611S). The Gibson reaction was then transformed into Stbl2 cells, and a small aliquot of the transformation culture was plated to ensure a transformation efficiency of >10X the library diversity. Library assembly and diversity was confirmed by sequencing on an Illumina MiSeq.

**Mammalian display cell culture protocol**

All 293t cells were grown in high glucose DMEM supplemented with Glutamax and pyruvate, 10% fetal bovine serum, and 1% penicillin/streptomycin. Cells were transduced at an MOI of approximately 0.1. 24h after transduction, 2μg/ml of puromycin was added to the cells, and cells were constantly grown in media containing 2μg/ml puromycin for subsequent steps. Once a sufficient number of cells had grown, approximately 10 million cells were physically dislodged from the cell culture flask. They were then washed in PBS, followed by staining with monoclonal antibodies of interest for 15 minutes at RT. Cells were then washed again with PBS, following which they were stained with anti-Myc FITC (Invitrogen, 13-2511) and anti-human IgG BV421 (BD Biosciences, 562581) antibodies for 15 minutes at RT. Cells were washed twice, resuspended in PBS, and finally stained with 7-AAD (ThermoFisher, A1310) before sorting. Cells were collected by sorting on a BD FACS Melody sorter. Cells were gated on singlets, 7-AAD-, and Myc+, following which the top 1-5% of iGL binding cells were collected in DMEM with 50% FBS. In addition, control cells that were unstained with any iGL and only Myc+ were sorted.

**Mammalian display sequencing**

Genomic DNA was extracted from sorted cells using a GenElute mammalian genomic DNA miniprep kit (Millipore Sigma, G1N70) according to the manufacturers instructions. Forward and reverse primers containing the Illumina P5 and P7 adapters respectively were designed to target regions of the Q23.PDGFR construct outside the area of the scanning NNK library, and used to amplify the region of interest. Amplified libraries were then purified using AMPureXP beads, and checked on a BioAnalyzer instrument to confirm that amplicons were of the correct size. The libraries were then sequenced on an Illumina MiSeq instrument using a 600 cycle MiSeq v3 kit.

**Mammalian display data analysis**
Following sequencing, reads were adaptor trimmed and paired. Library mutations in each sequence were identified by the presence of synonymous barcoding mutations flanking each NNK residue. The total number of cells bearing each mutation across the length of the NNK library was determined, and amino acid frequencies were calculated by dividing the number of cells with a particular amino acid mutation by the total number of cells bearing any mutation at that residue. Relative enrichments of mutations in the libraries were calculated via the ratio of the frequency of a sequence in the sorted library over the frequency of the same sequence in the control cells. Enrichments resulting from a low number of mutation counts in the control library were discarded.

**SHIV infection of rhesus macaques**
All Indian rhesus macaques were housed at Bioqual Inc (Rockville MD), according to AAALAC standards. All experiments were approved by the University of Pennsylvania and Bioqual Institutional Animal Care and Use Committees. Three days prior to SHIV infection, rhesus macaques were subcutaneously injected with 25mg/kg anti-CD8α mAb (MT807R1). Macaques were inoculated with SHIV by intravenous infusion. Blood draws, processing, and storage was performed as previously described (21).

**Env single genome sequencing**
3’ SHIV half genomes were sequenced as previously described (22). Briefly, viral RNA was synthesized from plasma virions using the Qiagen BioRobot EZ1 Workstation with EZ1 Virus Mini Kit v2.0 (Qiagen). Viral RNA was then used to synthesize cDNA using SuperScript III reverse transcriptase (Invitrogen). cDNA was then serially diluted in 96 well plates and amplified using nested PCR such that <30% of wells were PCR-positive. Positive wells were sequenced using an Illumina Miseq sequencer. Sequences from wells that showed multiple unique Env sequences were excluded. All Env sequences reported in this study are available at Genbank under accession numbers XXXX.

### Figures

A

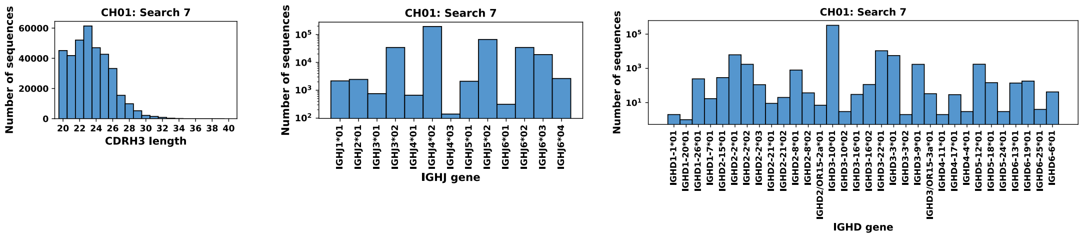


B

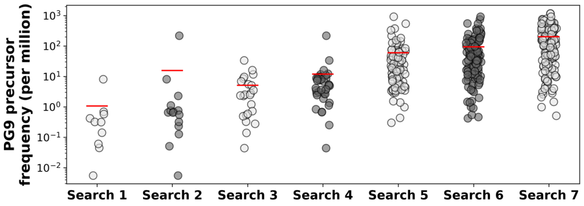


| Search | Motif register | JH gene | VH gene | # donors | # donors with 1M+ sequences |
|---|---|---|---|---|---|
| 1 | +/- 0 | J6 | V3 | 10/215 | 9/37 |
| 2 | +/- 0 | Any | V3 | 15/215 | 12/37 |
| 3 | +/- 2 | J6 | V3 | 23/215 | 16/37 |
| 4 | +/- 2 | Any | V3 | 37/215 | 21/37 |
| 5 | +/- 4 | J6 | V3 | 99/215 | 30/37 |
| 6 | +/- 4 | Any | V3 | 123/215 | 34/37 |
| 7 | +/- 4 | Any | Any | 152/215 | 35/37 |

C

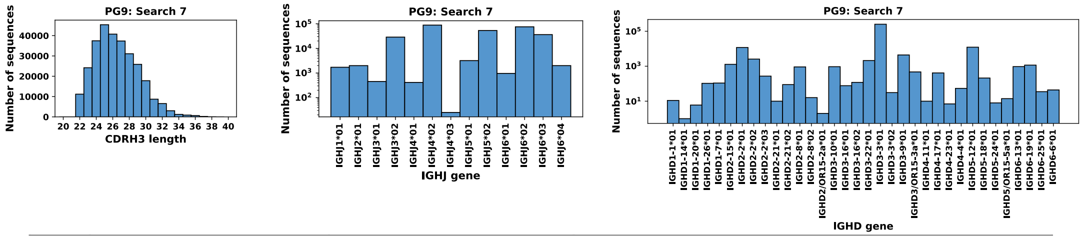


D

| Search | Regex | Description | All CDRH3 lengths | | | | |
|---|---|---|---|---|---|---|---|
| | | | From 215 donors (full set) | | | From 1M+ seq. donors | |
| | | | # donors | # sequences | Avg. freq. | # donors | # sequences |
| 1 | **'^.{2,10}EDDYGYYYT.{6,20}$'** | Conserved unmutated NHP D gene | 2/215 | 20 | 6.37 | 1/37 | 19 |
| 2 | **'^.{2,}EDDYG.{6,}$'** | D gene core motif (rhesus) | 109/215 | 92953 | 97.09 | 33/37 | 91717 |
| 3 | **'^.{2,}[EG]DDYG.{6,}$'** | D gene core motif (V033) | 155/215 | 317625 | 249.09 | 36/37 | 311198 |

| Search | Regex | Description | CDRH3 lengths >= 20 | | | | |
|---|---|---|---|---|---|---|---|
| | | | From 215 donors (full set) | | | From 1M+ seq. donors | |
| | | | # donors | # sequences | Avg. freq. | # donors | # sequences |
| 1 | **'^.{2,10}EDDYGYYYT.{6,20}$'** | Conserved unmutated NHP D gene | 2/215 | 20 | 6.37 | 1/37 | 19 |
| 2 | **'^.{2,}EDDYG.{6,}$'** | D gene core motif (rhesus) | 55/215 | 9728 | 19.87 | 24/37 | 9572 |
| 3 | **'^.{2,}[EG]DDYG.{6,}$'** | D gene core motif (V033) | 87/215 | 49922 | 44.46 | 31/37 | 49299 |

| Search | Regex | Description | CDRH3 lengths >= 23 | | | | |
|---|---|---|---|---|---|---|---|
| | | | From 215 donors (full set) | | | From 1M+ seq. donors | |
| | | | # donors | # sequences | Avg. freq. | # donors | # sequences |
| 1 | **'^.{2,10}EDDYGYYYT.{6,20}$'** | Conserved unmutated NHP D gene | 1/215 | 5 | 2.71 | 1/37 | 5 |
| 2 | **'^.{2,}EDDYG.{6,}$'** | D gene core motif (rhesus) | 32/215 | 3708 | 20.50 | 21/37 | 3663 |
| 3 | **'^.{2,}[EG]DDYG.{6,}$'** | D gene core motif (V033) | 58/215 | 12023 | 21.45 | 27/37 | 11855 |

**Fig. S1. Diverse CDRH3 sequences were found in OAS database searches. A.** CH01 Search 7 CDRH3 length distribution, the distribution of IGHJ genes, and the distribution of IGHD genes, before removal of redundant sequences. **B.** PG9-like antibody search results from the OAS database. Each point represents a donor that had at least one sequence matching the search criteria, as detailed in the table. **C.** PG9 Search 7 CDRH3 length distribution, the distribution of IGHJ genes, and the distribution of IGHD genes, before removal of redundant sequences. **D.** Results of the rhesus-related OAS searches broken down by CDRH3 length.

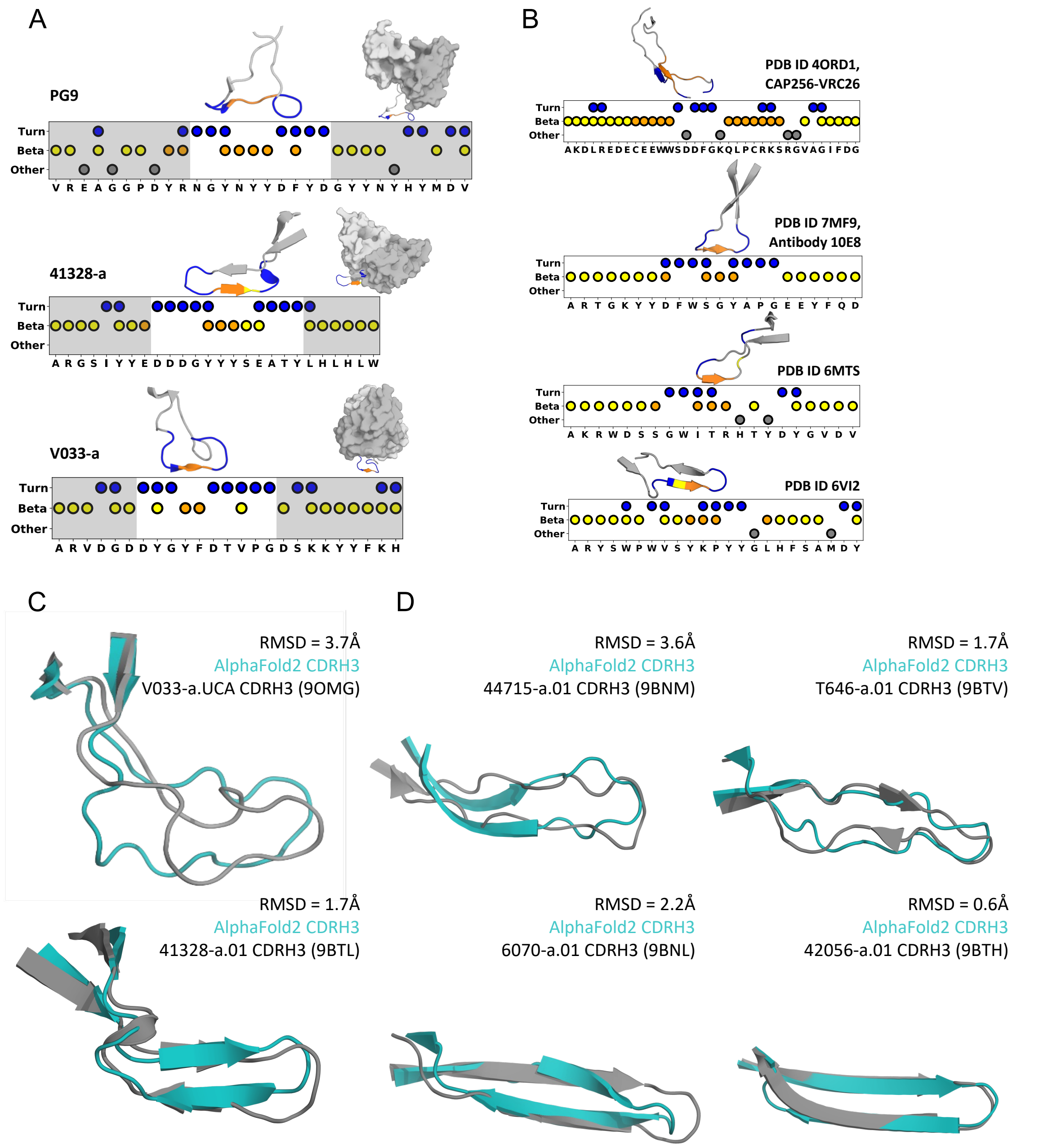


**Fig. S2. Structural classification and prediction of axe-like folds. A.** Our structural definition of an axe-like CDRH3 applied to PG9 (PDB ID: 5VJ6), 41328-a (PDB ID: 9BTL), and V033-a (PDB ID: 9BNP), with each respective CDRH3 shown above each scatter plot, as well as a structure of their variable heavy (gray) and light chains (light gray) shown as a surface representation with the CDRH3 pulled out. **B.** CDRH3 residue classifications (scatter plots) and CDRH3 structures for the antibodies classified as axe-like from the SAbDab database (excluding CH01-lineage structures, the second CAP256-VRC26 structure (PDB ID: 4ORG), and the second 10E8 structure (PDB ID: 5JNY). **C-D**, Structural alignments of AlphaFold2 predictions of rhesus V2 apex bnAb CDRH3s with their corresponding Cryo-EM structures for axe-like (C) and needle-like (D) lineages. AlphaFold predictions are shown in teal, and Cryo-EM structures are shown in gray.

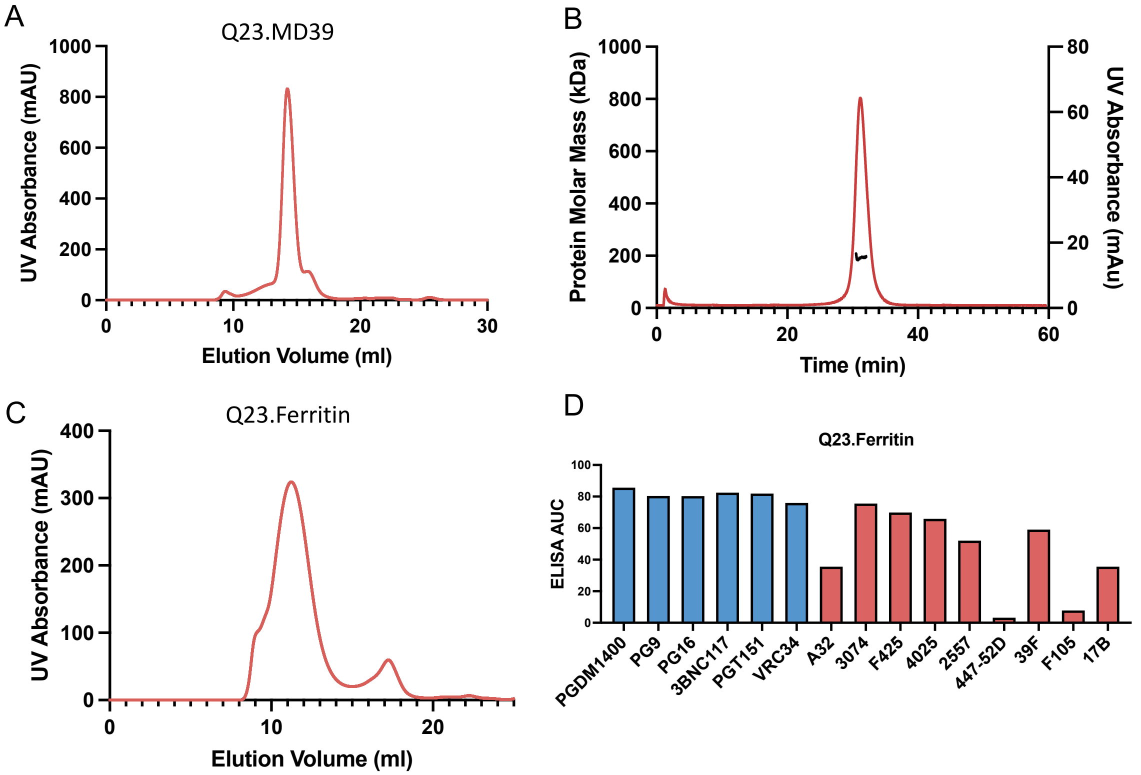


**Fig. S3. Characterization of Q23-based immunogens. A.** Size-exclusion chromatography trace of Q23.MD39. **B.** Size exclusion chromatography with multi-angle light scattering trace of Q23.MD39. Red line indicates UV absorbance. Black line indicates protein molecular weight of the main peak. **C.** Size-exclusion chromatography trace of Q23.Ferritin. **D.** Antigenic profile of Q23.Ferritin against a panel of broadly neutralizing and non-neutralizing antibodies.

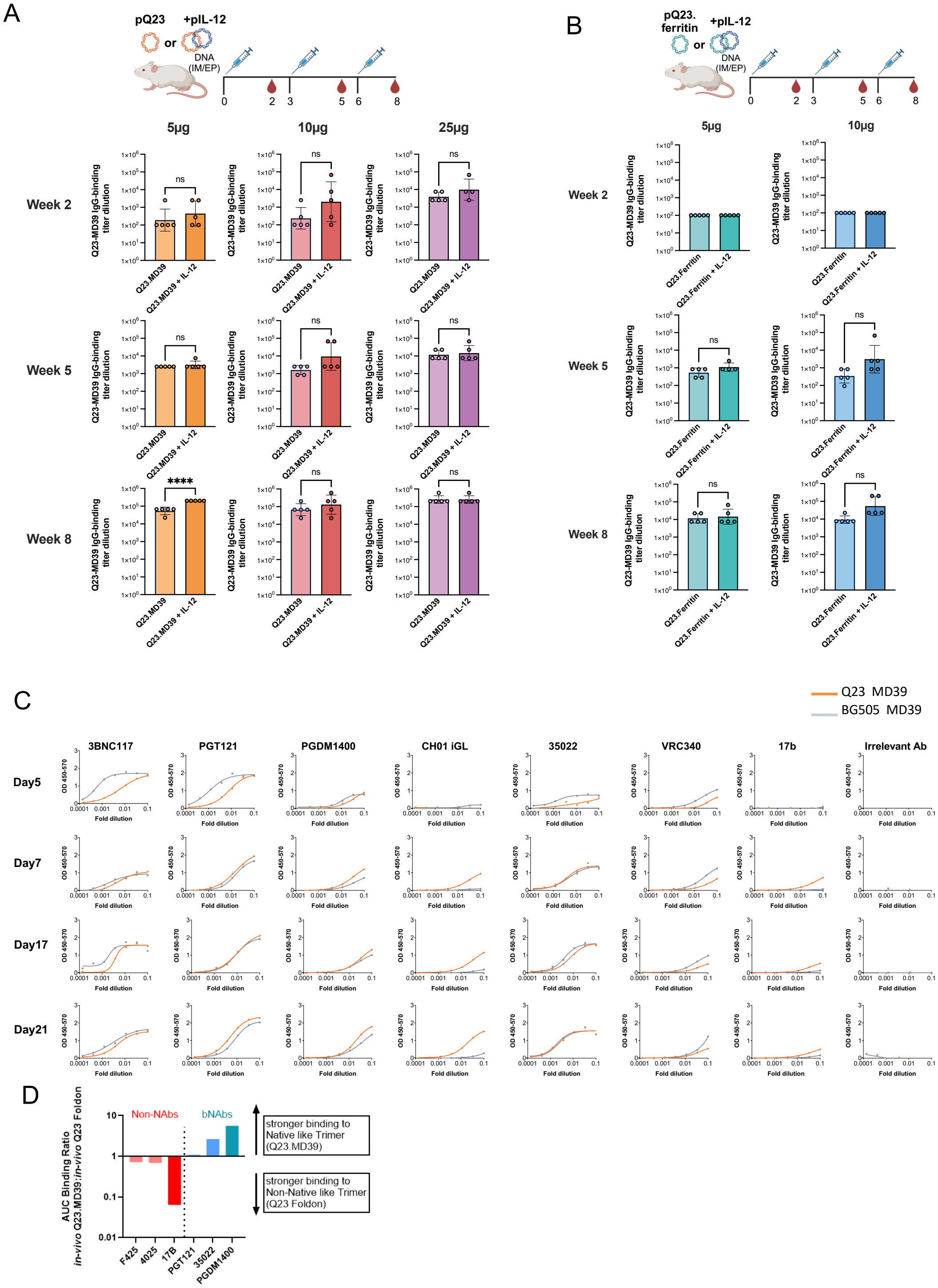

**Fig. S4. Immunogenicity of DNA-delivered Q23 immunogens.** Time course of vaccine-specific serum-IgG binding endpoint titers in WT BALB/c mice immunized with predetermined dosages of (**A**) 5ug, 10ug and 25μg of DNA-delivered Q23.MD39 and (**B**) 5ug and 10ug of DNA delivered Q23.Ferritin. Circles indicate individual mice. **C.** Antigen Conformation Tracing In Vivo by ELISA (ACTIVE) comparison or DNA-delivered, *in vivo* produced Q23.MD39 or BG505.MD39 against a panel of bnAbs and non-nAbs. **D.** Ratio of ELISA area under the curve binding of pQ23.MD39 to pQ23-foldon in an ACTIV assay against a panel of HIV bnAbs and non-nAbs.

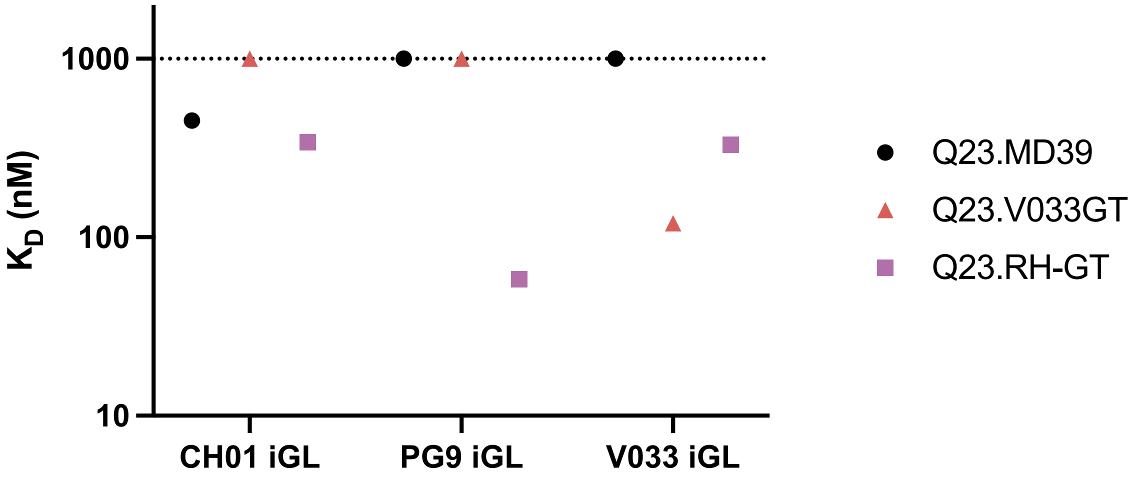


**Fig. S5. SPR binding affinities of Q23.MD39, Q23.V033GT, and Q23.RH-GT to axe-like bnAb iGLs.**

A

| | TZM-bl Titer (IU/µl) | p27 ng/ml |
|---|---|---|
| **SHIV-Q23.17** | 1844 | 541 |
| **SHIV-Q23.V033GT** | 2594 | 658 |

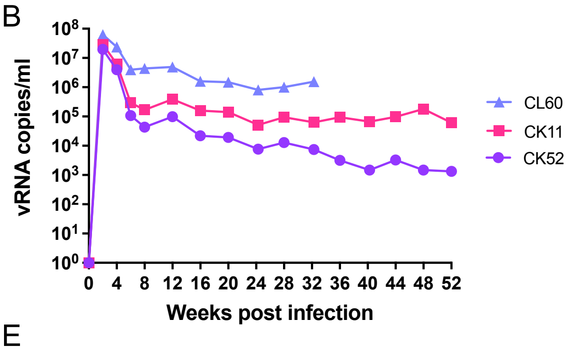


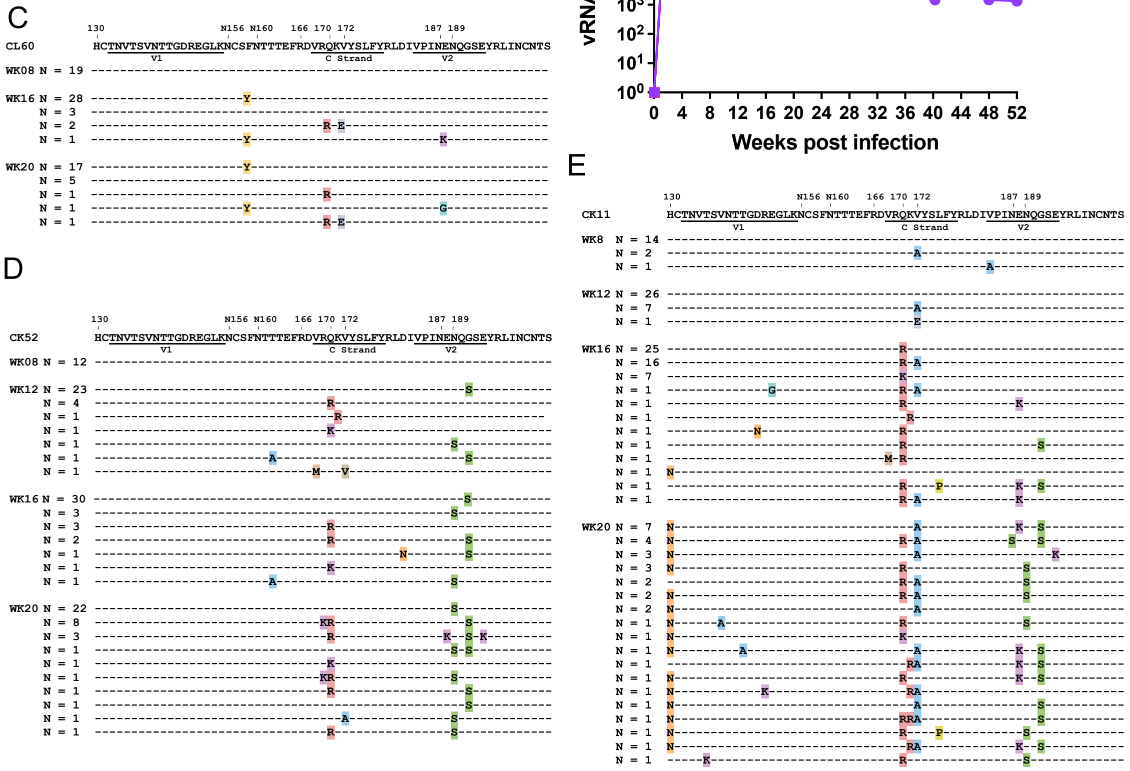


**Fig. S6. Infectivity, replication kinetics, and evolution of SHIV-Q23.V033GT. A.** TZM-bl and p27 antigen ELISA titers of WT SHIV-Q23.17 and SHIV-Q23.V033GT. **B.** Viral replication kinetics in SHIV-Q23.V033GT infected macaques. Macaque CL60 was euthanized 32 weeks post-infection due to symptoms associated with rapid progression to AIDS. **C-E.** Single genome sequencing of the V1V2 region (HXB2 residues 130-199) of circulating plasma virus in macaques CL60 (C), CK52 (D), and CK11 (E). Mismatches to the transmitted/founder SHIV are highlighted and colored by amino acid identity. N = number of viruses. Numbers over the reference sequence indicate HXB2 residue numbers, and timepoints of virus isolation are labelled on the left of the topmost sequence.

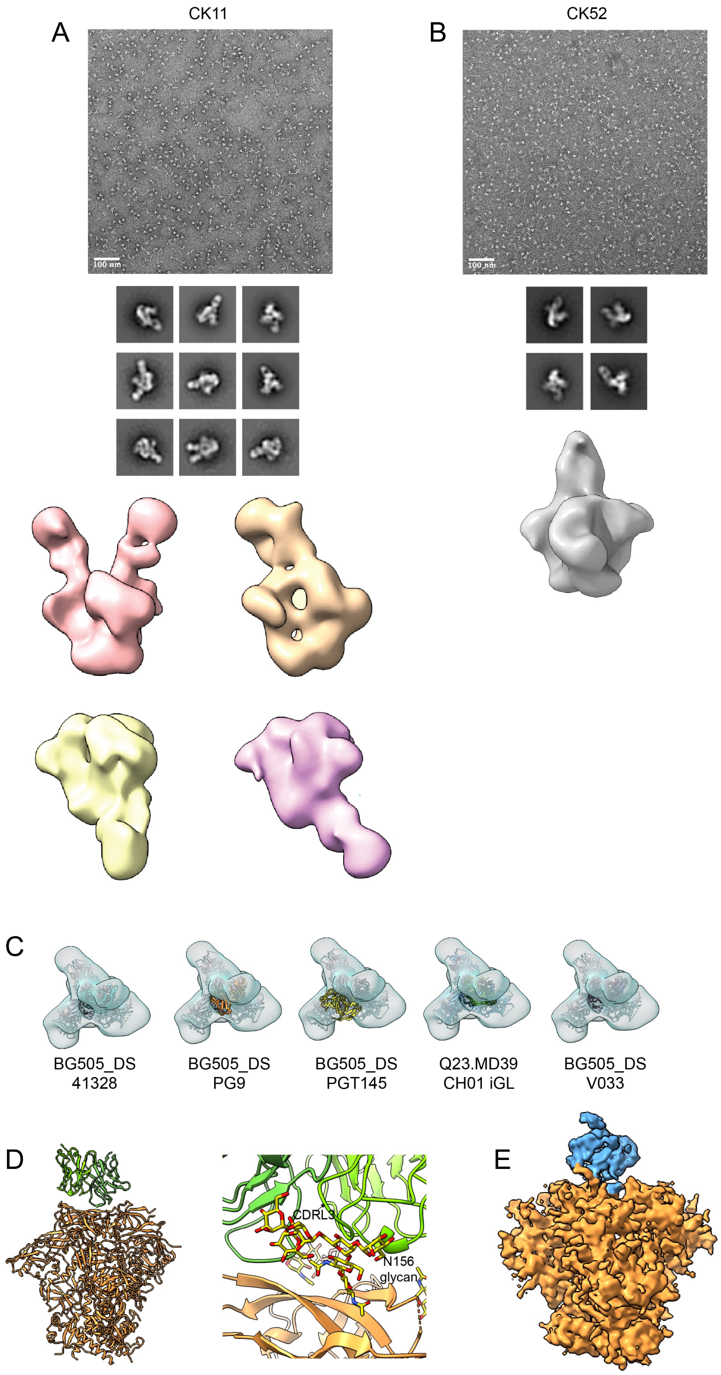


**Fig. S7. NS-EMPEM of the polyclonal antibody responses to Q23.V033GT in rhesus macaques CK11 and CK52. A.** Representative raw micrograph, 2D classes, and 3D reconstructions of Fab/Q23.V033GT complexes from CK11 at week 24 post-infection. **B.** Representative raw micrograph, 2D classes, and 3D reconstructions of Fab/Q23.V033GT complexes from CK52 at week 24 post-infection. **C.** 3D reconstruction of a NS-EM V2 apex binding class from CK52 at week 24, docked with atomic models of BG505_DS/41328, BG505_DS/V033, BG505_DS/PG9 (PDB: 8FL1), BG505 SOSIP/PGT145 (PDB: 8FLW) and Q23.MD39/CH01 iGL (PDB: XXXX). **D.** Atomic model of CK52.1 in complex with Q23.V033GT and a zoomed view of the CK52.1 in engaging with the N160 glycan. **E.** CryoEM density map of a V2 apex antibody binding to the Q23.V033GT from the CK11 polyclonal serum.

```
                                                                                        106
Env Q23.17   AVENLWVTVYYGVPVWRDADTTLFCASDAKAYETEKHNVWATHACVPTDPNPQEIHLDNVTEKFNMWKNNMVEQMHTDIISLWDQSLKPCVKLTPLCVTL
Q23.MD39     AVENLWVTVYYGVPVWRDADTTLFCASDAKAYETEKHNVWATHACVPTDPNPQEIHLDNVTEKFNMWKNNMVEQMHEDIISLWDQSLKPCVKLTPLCVTL
Q23.KO       AVENLWVTVYYGVPVWRDADTTLFCASDAKAYETEKHNVWATHACVPTDPNPQEIHLDNVTEKFNMWKNNMVEQMHEDIISLWDQSLKPCVKLTPLCVTL
Q23.V033GT   AVENLWVTVYYGVPVWRDADTTLFCASDAKAYETEKHNVWATHACVPTDPNPQEIHLDNVTEKFNMWKNNMVEQMHEDIISLWDQSLAPCVKLTPLCVTL
Q23.RH-GT    AVENLWVTVYYGVPVWRDADTTLFCASDAKAYETEKHNVWATHACVPTDPNPQEIHLDNVTEKFNMWKNNMVEQMHEDIISLWDQSLKPCVKLTPLCVTL
Q23.Ferritin AVENLWVTVYYGVPVWRDADTTLFCASDAKAYETEKHNVWATHACVPTDPNPQEIHLDNVTEKFNMWKNNMVEQMHEDIISLWDQSLKPCVKLTPLCVTL
Q23.PDGFR    AVENLWVTVYYGVPVWRDADTTLFCASDAKAYETEKHNVWATHACVPTDPNPQEIHLDNVTEKFNMWKNNMVEQMHEDIISLWDQSLKPCVKLTPLCVTL

             130               156 160  165  170
Env Q23.17   HCTNVTSVNTTGDREGLKNCSFNMTTELRDKRQKVYSLFYRLDIVPINENQGSEYRLINCNTSAITQACPKVSFEPIPIHYCTPAGFAILKCKDEGFNGT
Q23.MD39     HCTNVTSVNTTGDREGLKNCSFNMTTELRDKRQKVYSLFYRLDIVPINENQGSEYRLINCNTSAITQACPKVSFEPIPIHYCTPAGFAILKCKDEGFNGT
Q23.KO       HCTNVTSVNTTGDREGLKNCSFNMTTELRDKEQEVYSLFYRLDIVPINENQGSEYRLINCNTSAITQACPKVSFEPIPIHYCTPAGFAILKCKDEGFNGT
Q23.V033GT   HCTNVTSVNTTGDREGLKNCSFNTTTEFRDVRQKVYSLFYRLDIVPINENQGSEYRLINCNTSAITQACPKVSFEPIPIHYCTPAGFAILKCKDEGFNGT
Q23.RH-GT    HCTNVTSVQTTGDREGLKNCSFNSTTELRDKRQKVYSLFKRLDIVPINENQGSEYRLINCNTSAITQACPKVSFEPIPIHYCTPAGFAILKCKDEGFNGT
Q23.Ferritin HCTNVTSVNTTGDREGLKNCSFNMTTELRDKRQKVYSLFYRLDIVPINENQGSEYRLINCNTSAITQACPKVSFEPIPIHYCTPAGFAILKCKDEGFNGT
Q23.PDGFR    HCTNVTSVNTTGDREGLKNCSFNMTTELRDKRQKVYSLFYRLDIVPINENQGSEYRLINCNTSAITQACPKVSFEPIPIHYCTPAGFAILKCKDEGFNGT

Env Q23.17   GLCKNVSTVQCTHGIKPVVSTQLLLNGSLAEKNITIRSENITNNAKIIIVQLVQPVTIKCIRPNNNTRKSIRIGPGQAFYATGDIIGDIRQAHCNVTRSR
Q23.MD39     GLCKNVSTVQCTHGIKPVVSTQLLLNGSLAEKNIIIRSENITNNAKIIIVQLVQPVTIKCIRPNNNTVKSIRIGPGQAFYYTGDIIGDIRQAHCNVTRSR
Q23.KO       GLCKNVSTVQCTHGIKPVVSTQLLLNGSLAEKNIIIRSENITNNAKIIIVQLVQPVTIKCIRPNNNTVKSIRIGPGQAFYYTGDIIGDIRQAHCNVTRSR
Q23.V033GT   GLCKNVSTVQCTHGIKPVVSTQLLLNGSLAEKNIIIRSENITNNAKIIIVQLVQPVTIKCIRPNNNTVKSIRIGPGQAFYYTGDIIGDIRQAHCNVTRSR
Q23.RH-GT    GLCKNVSTVQCTHGIKPVVSTQLLLNGSLAEKNIIIRSENITNNAKIIIVQLVQPVTIKCIRPNNNTVKSIRIGPGQAFYYTGDIIGDIRQAHCNVTRSR
Q23.Ferritin GLCKNVSTVQCTHGIKPVVSTQLLLNGSLAEKNIIIRSENITNNAKIIIVQLVQPVTIKCIRPNNNTVKSIRIGPGQAFYYTGDIIGDIRQAHCNVTRSR
Q23.PDGFR    GLCKNVSTVQCTHGIKPVVSTQLLLNGSLAEKNIIIRSENITNNAKIIIVQLVQPVTIKCIRPNNNTVKSIRIGPGQAFYYTGDIIGDIRQAHCNVTRSR

Env Q23.17   WNKTLQEVAEKLRTYFGNKTIIFANSSGGDLEITTHSFNCGGEFFYCNTSGLFNSTWYVNSTWNDTDSTQESNDTITLPCRIKQIINMWQRAGQAMYAPP
Q23.MD39     WNKTLQEVAEKLRTYFGNKTIIFAQSSGGDLEITTHSFNCGGEFFYCNTSGLFNSTWYVNSTWNDTDSTQESNDTITLPCRIKQIINMWQRAGQAMYAPP
Q23.KO       WNKTLQEVAEKLRTYFGNKTIIFAQSSGGDLEITTHSFNCGGEFFYCNTSGLFNSTWYVNSTWNDTDSTQESNDTITLPCRIKQIINMWQRAGQAMYAPP
Q23.V033GT   WNKTLQEVAEKLRTYFGNKTIIFAQSSGGDLEITTHSFNCGGEFFYCNTSGLFNSTWYVNSTWNDTDSTQESNDTITLPCRIKQIINMWQRAGQAMYAPP
Q23.RH-GT    WNKTLQEVAEKLRTYFGNKTIIFAQSSGGDLEITTHSFNCGGEFFYCNTSGLFNSTWYVNSTWNDTDSTQESNDTITLPCRIKQIINMWQRAGQAMYAPP
Q23.Ferritin WNKTLQEVAEKLRTYFGNKTIIFAQSSGGDLEITTHSFNCGGEFFYCNTSGLFNSTWYVNSTWNDTDSTQESNDTITLPCRIKQIINMWQRAGQAMYAPP
Q23.PDGFR    WNKTLQEVAEKLRTYFGNKTIIFAQSSGGDLEITTHSFNCGGEFFYCNTSGLFNSTWYVNSTWNDTDSTQESNDTITLPCRIKQIINMWQRAGQAMYAPP

Env Q23.17   IPGVIKCESNITGLLLTRDGGKDNNVNETFRPGGGDMRDNWRSELYKYKVVEIEPLGVAPTRAKRRVVE--REKRAVGIGAVFLGFLGAAGSTMGATSIT
Q23.MD39     IPGVIKCESNITGLLLTRDGGKDNNVNETFRPGGSDMRDNWRSELYKYKVVEIEPLGVAPTRCKRRVVERRRRRRAVGIGAVSLGFLGAAGSTMGAASIT
Q23.KO       IPGVIKCESNITGLLLTRDGGKDNNVNETFRPGGSDMRDNWRSELYKYKVVEIEPLGVAPTRCKRRVVERRRRRRAVGIGAVSLGFLGAAGSTMGAASIT
Q23.V033GT   IPGVIKCESNITGLLLTRDGGKDNNVNETFRPGGSDMRDNWRSELYKYKVVEIEPLGVAPTRCKRRVVERRRRRRAVGIGAVSLGFLGAAGSTMGAASIT
Q23.RH-GT    IPGVIKCESNITGLLLTRDGGKDNNVNETFRPGGSDMRDNWRSELYKYKVVEIEPLGVAPTRCKRRVVERRRRRRAVGIGAVSLGFLGAAGSTMGAASIT
Q23.Ferritin IPGVIKCESNITGLLLTRDGGKDNNVNETFRPGGSDMRDNWRSELYKYKVVEIEPLGVAPTRCKRRVVERRRRRRAVGIGAVSLGFLGAAGSTMGAASIT
Q23.PDGFR    IPGVIKCESNITGLLLTRDGGKDNNVNETFRPGGSDMRDNWRSELYKYKVVEIEPLGVAPTRCKRRVVERRRRRRAVGIGAVSLGFLGAAGSTMGAASIT

Env Q23.17   LTVQARQLLSGIVQQQNNLLRAIEAQQHLLKLTVWGIKQLQARVLAVERYLRDQQLLGIWGCSGKLICTTNVPWNSSWSNKSLDEIWNNMTWLQWDKEIN
Q23.MD39     LTVQARQLLSGIVQQQNNLLRAPEPQQHLLKDTHWGIKQLQARVLAVEHYLRDQQLLGIWGCSGKLICCTNVPWNSSWSNKSLDEIWNNMTWLQWDKEIN
Q23.KO       LTVQARQLLSGIVQQQNNLLRAPEPQQHLLKDTHWGIKQLQARVLAVEHYLRDQQLLGIWGCSGKLICCTNVPWNSSWSNKSLDEIWNNMTWLQWDKEIN
Q23.V033GT   LTVQARQLLSGIVQQQNNLLRAPEPQQHLLKDTHWGIKQLQARVLAVEHYLRDQQLLGIWGCSGKLICCTNVPWNSSWSNKSLDEIWNNMTWLQWDKEIN
Q23.RH-GT    LTVQARQLLSGIVQQQNNLLRAPEPQQHLLKDTHWGIKQLQARVLAVEHYLRDQQLLGIWGCSGKLICCTNVPWNSSWSNKSLDEIWNNMTWLQWDKEIN
Q23.Ferritin LTVQARQLLSGIVQQQNNLLRAPEPQQHLLKDTHWGIKQLQARVLAVEHYLRDQQLLGIWGCSGKLICCTNVPWNSSWSNKSLDEIWNNMTWLQWDKEIN
Q23.PDGFR    LTVQARQLLSGIVQQQNNLLRAPEPQQHLLKDTHWGIKQLQARVLAVEHYLRDQQLLGIWGCSGKLICCTNVPWNSSWSNKSLDEIWNNMTWLQWDKEIN

Env Q23.17   NYTQLIYRLIEESQNQQEKNEKELLELDKWANLWSWFDISNWLWYIKIFIIIVGGLIGLRIVFAVLSVINRVRQGYSPLSFQTHTPNPRGLDRPERIEEE
Q23.MD39     NYTQLIYRLIEESQNQQEKNEKELLELD------------------------------------------------------------------------
Q23.KO       NYTQLIYRLIEESQNQQEKNEKELLELD------------------------------------------------------------------------
Q23.V033GT   NYTQLIYRLIEESQNQQEKNEKELLELD------------------------------------------------------------------------
Q23.RH-GT    NYTQLIYRLIEESQNQQEKNEKELLELD------------------------------------------------------------------------
Q23.Ferritin NYTQLIYRLIEESQNQQEKNEKELLELDGSGGLSKDIIKLLNEQVNKEMQSSNLYMSMSSWCYTHSLDGAGLFLFDHAAEEYEHAKKLIIFLNENNVPVQ
Q23.PDGFR    NYTQLIYRLIEESQNQQEKNEKELLELDGGSGGAGGSEQKLISEEDLGGSGGAGGSAVGQDTQEVIVVPHSLPFKVVVISAILALVVLTIISLIILIMLW

Env Q23.17   DGEQGRGRSIRLVSGFLALAWDDLRSLCLFSYHRLRDFILIAARTVELLGHSSLKGLRLGWEGIKYLWNLLSYW-GRELKISAINLVDTIAIAVAGWTDR
Q23.MD39     ----------------------------------------------------------------------------------------------------
Q23.KO       ----------------------------------------------------------------------------------------------------
Q23.V033GT   ----------------------------------------------------------------------------------------------------
Q23.RH-GT    ----------------------------------------------------------------------------------------------------
Q23.Ferritin LTSISAPEHKFEGLTQIFQKAYEHEQHISESINNIVDHAIKSKDHATFNFLQWYVAEQHEEEVLFKDILDKIELIGNENHGLYLADQYVKGIAKSRKS--
Q23.PDGFR    QKKPR-----------------------------------------------------------------------------------------------

Env Q23.17   VIEIAQRIGRAILHIPVRIRQGLERALL
Q23.MD39     ----------------------------
Q23.KO       ----------------------------
Q23.V033GT   ----------------------------
Q23.RH-GT    ----------------------------
Q23.Ferritin ----------------------------
Q23.PDGFR    ----------------------------
```

**Fig. S8. Amino acid sequence alignment of Env Q23.17 and Q23.17-based constructs.** Mismatches to Q23.MD39 are highlighted. Numbers indicate HXB2 residue positions, and dashes indicate gaps in the alignment.

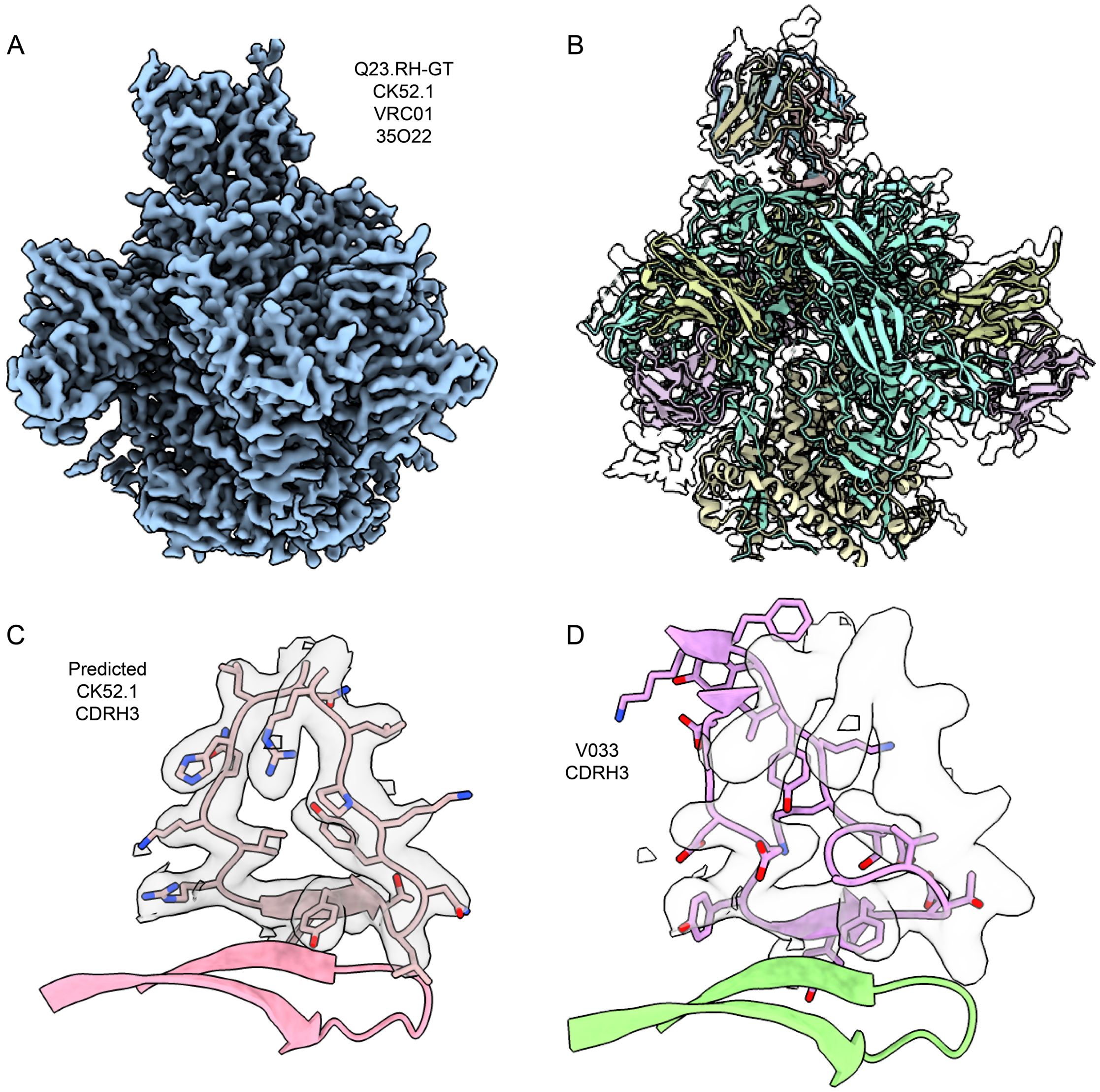


**Fig. S9. Cryo-EMPEM structure of Q23.RH-GT in complex with CK52.1, VRC01, and 35O22.** **A.** CryoEM density map of Q23.RH-GT in complex with CK52.1, VRC01, and 35O22. **B.** Docked models of Q23.RH-GT in complex with VRC01 and 35O22 along with the ModelAngelo predicted peptide fragments. **C.** Zoomed view of the predicted CK52.1 CDRH3 sequence aligned with the density map in A. **D.** The V033-a CDRH3 docked into the density map in A.

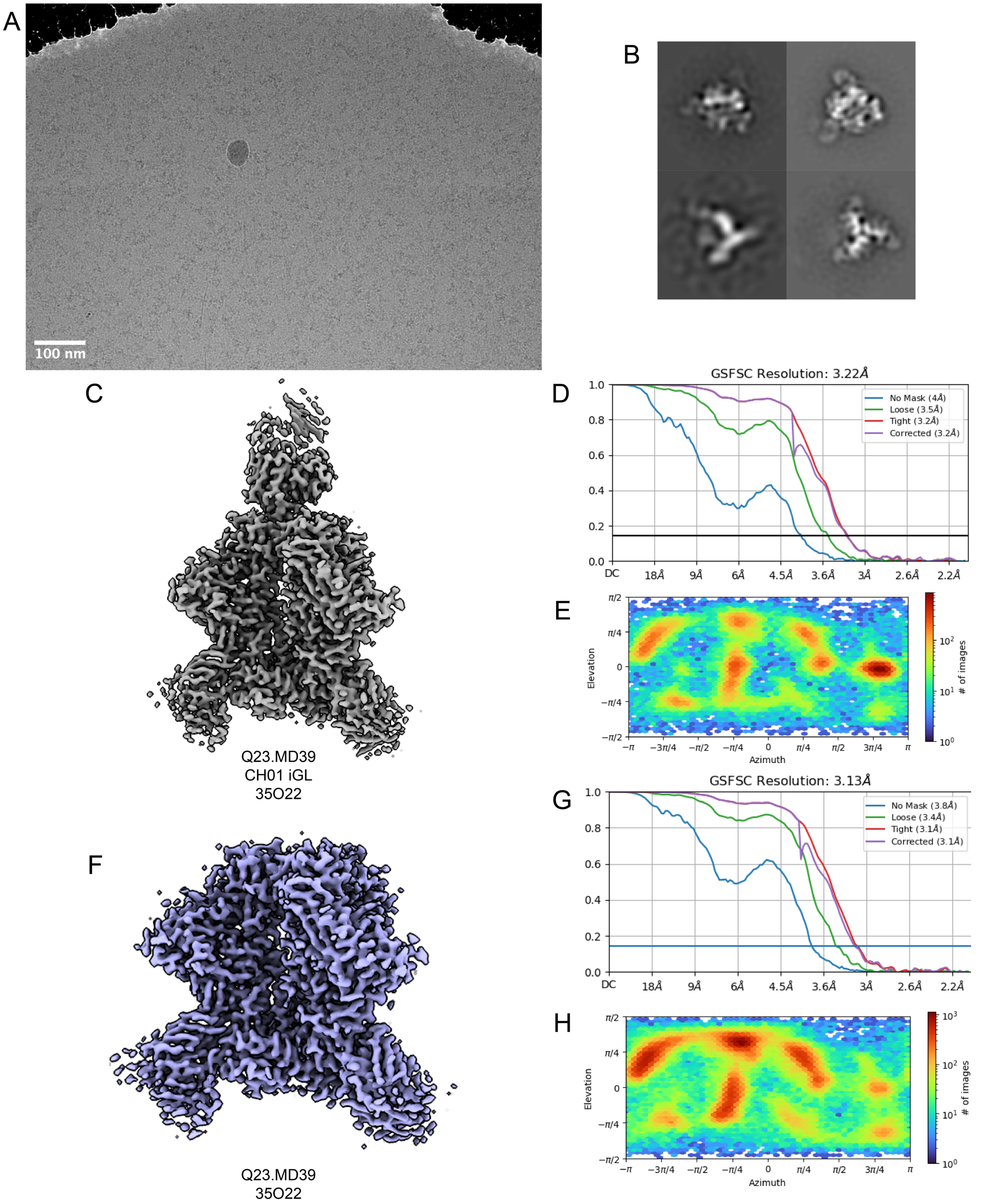


**Fig. S10. CryoEM details of Q23.MD39/35O22 with and without CH01 iGL. A**. A representative micrograph of the Q23.MD39/CH01iGL/35O22 dataset. **B**. Representative 2D Classes. **C**. CryoEM reconstruction of the Q23.MD39/35O22 in complex with CH01 iGL. **D-E** Gold standard Fourier shell correlation (GS-FSC) and particle orientation distribution of CryoEM reconstruction of Q23.MD39/35O22 in complex with CH01 iGL. **F**. CryoEM reconstruction of the Q23.MD39 with 35O22. (**G-H**) GS-FSC and particle orientation distribution of CryoEM reconstruction of Q23.MD39/35O22 in complex with CH01 iGL.

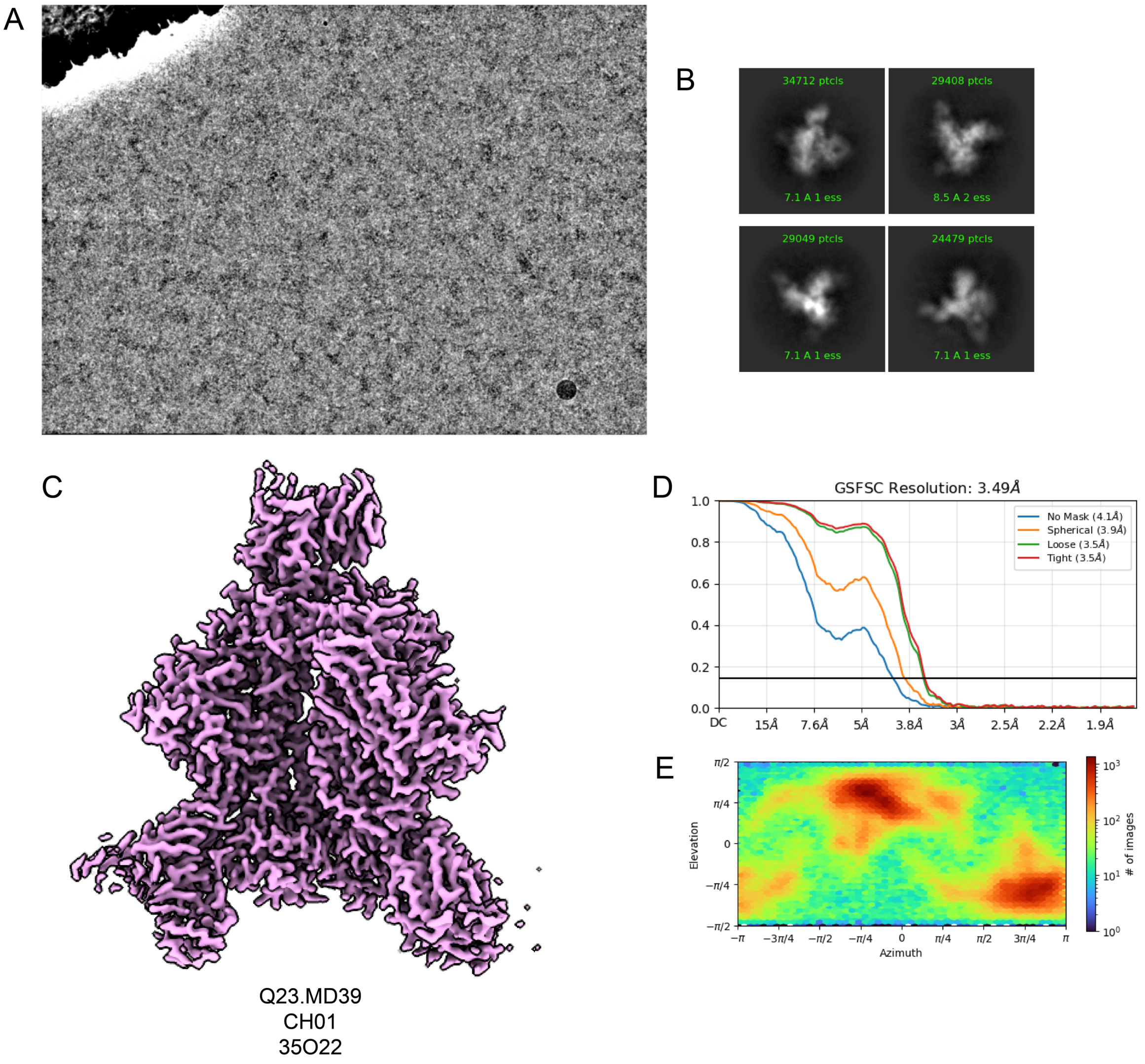


**Fig. S11. CryoEM details of Q23.MD39/35O22 with CH01. A**. A representative micrograph of the Q23.MD39/CH01/35O22 dataset. **B**. Representative 2D Classes. **C**. CryoEM reconstruction of the Q23.MD39/35O22 in complex with CH01. **D-E**. GS-FSC and particle orientation distribution of CryoEM reconstruction of Q23.MD39/35O22 in complex with CH01.

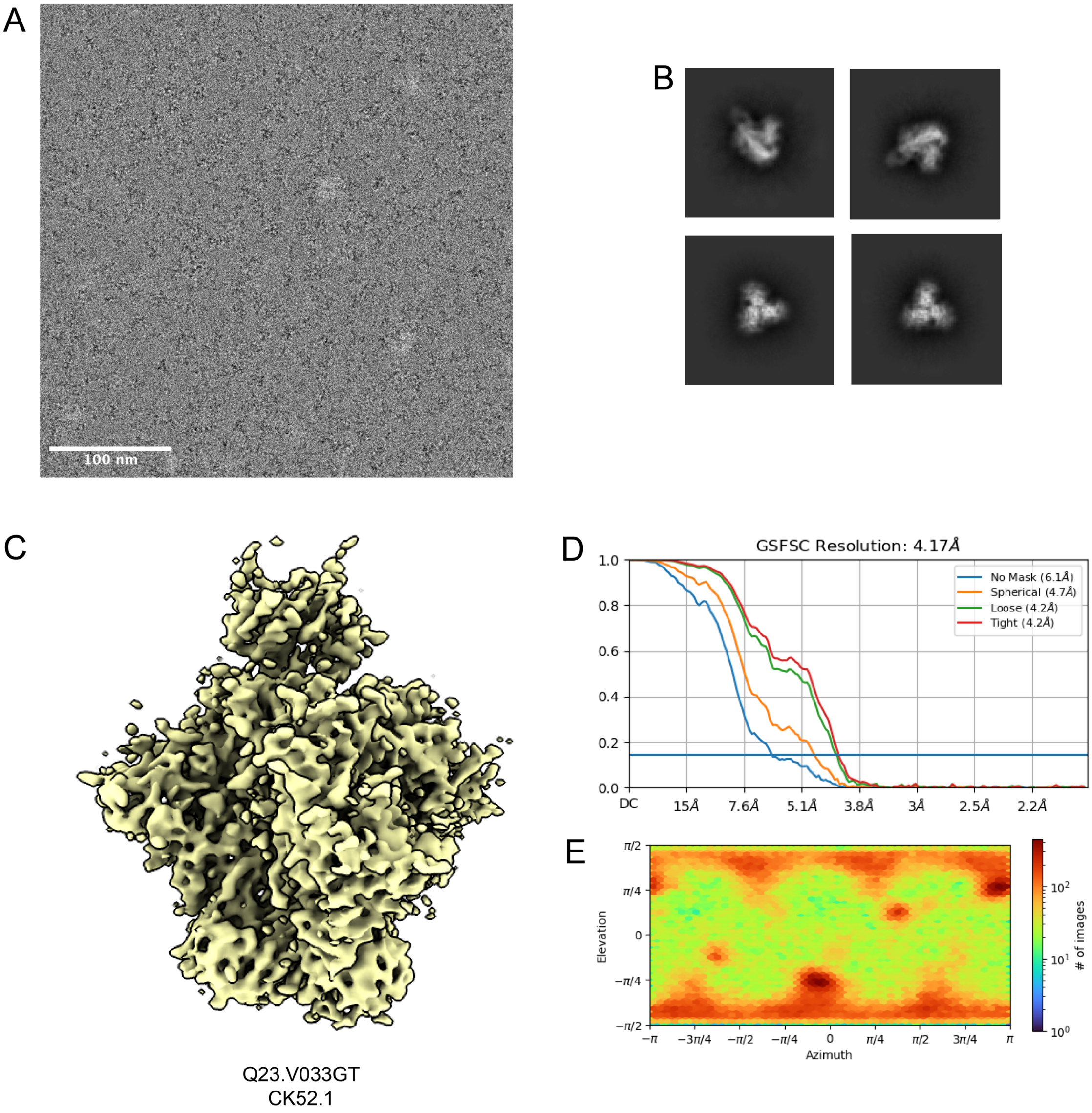


**Fig. S12. CryoEMPEM details of Q23.V033GT in complex with CK52 week 24 V2 apex antibody. A**. A representative micrograph of the Q23.V033GT in complex with CK52 week 24 polyclonal fabs. **B**. Representative 2D Classes. **C**. CryoEM reconstruction of the Q23.V033GT in complex with CK52.1. **D-E**. GS-FSC and particle orientation distribution of CryoEM reconstruction of Q23.V033GT in complex with CK52.1.

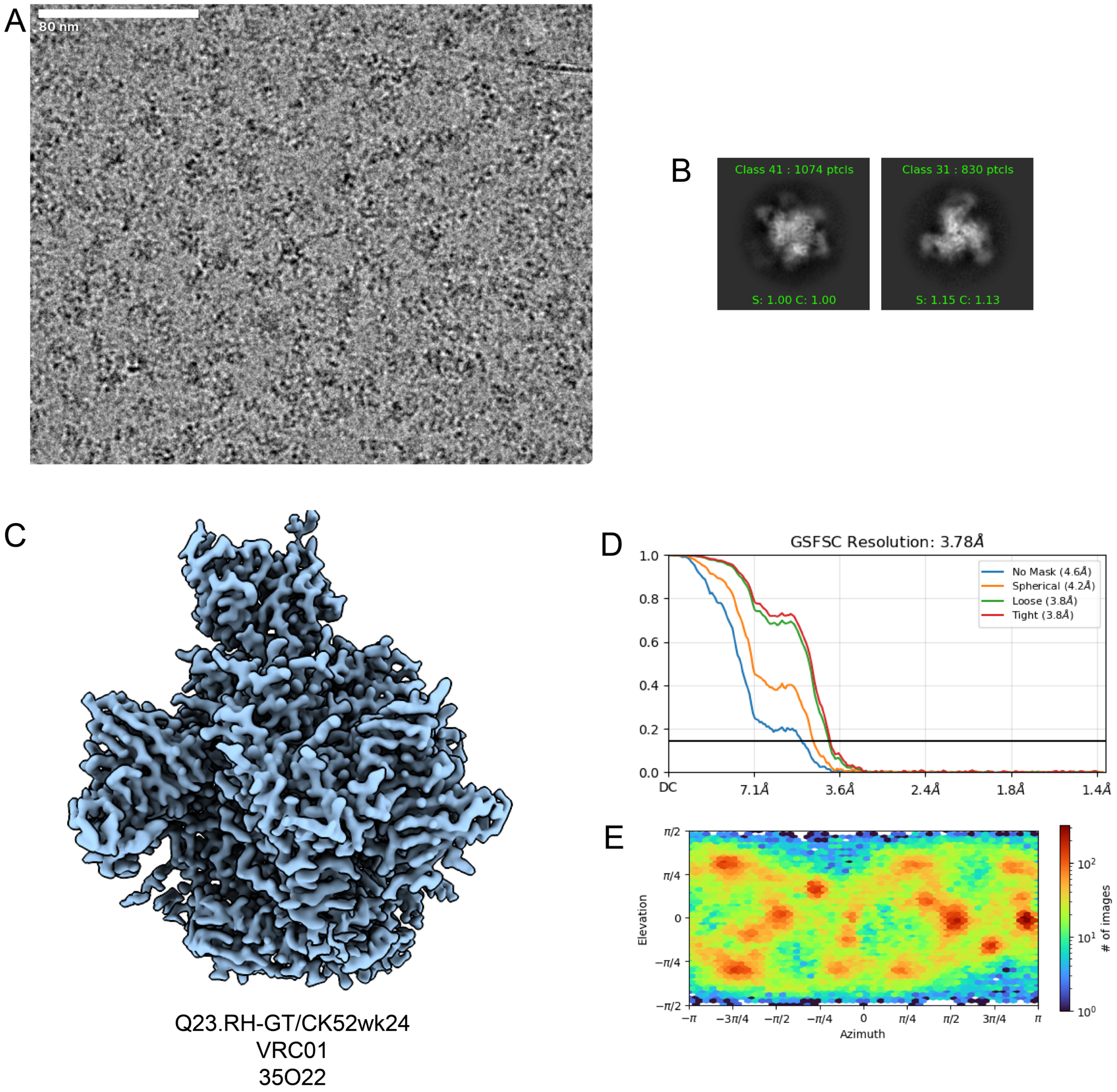


**Fig. S13. CryoEM details of Q23.RH-GT in complex with CK52 week 24 V2 apex antibody. A**. A representative micrograph of the Q23.RH-GT in complex with CK52 week 24 polyclonal fabs. **B**. Representative 2D Classes. **C**. CryoEM reconstruction of the Q23.RH-GT in complex with CK52.1. **D-E**. GS-FSC and particle orientation distribution of CryoEM reconstruction of Q23.RH-GT in complex with CK52.1.

### Tables

| Data Collection and Processing | Q23.MD39/CH01iGL / 35o22 | Q23.MD39/35O22 | Q23.MD39/CH01 35O22 | Q23.V033GT / CK52wk24 | Q23.RH-GT VRC01/35O22 CK52wk24 |
|---|---|---|---|---|---|
| Electron Microscope | Titan Krios | | Titan Krios | Glacios | Glacios 2 |
| Electron Detector | K3 Summit | | K3 Summit | Falcon 4 | Falcon 4i |
| Magnification | 81,000 | | 150,000 | 150,000 | 150,000 |
| Voltage (kV) | 300 | | 300 | 200 | 200 |
| Total Electron Exposure ($e^-$ /Å$^2$ ) | 40 | | 41 | 50 | 45 |
| Defocus Range (μm) | 1 - 2.2 | | 0.8-2.5 | 0.8-2.2 | 0.8-2.2 |
| Pixel Size (Å) | 1.05 | | 0.43 | 0.95 | 0.7 |
| Symmetry | C1 | C1 | C1 | C1 | C1 |
| Number of final particle images | 71,949 | 150,988 | 248,110 | 165,144 | 71,460 |
| Map Resolution (Å) | 3.22 | 3.13 | 3.49 | 4.17 | 3.78 |
| FSC Threshold | 0.143 | 0.143 | 0.143 | 0.143 | 0.143 |
| Sharpening Factor | 62.6 | 82.8 | 103 | 101.7 | 62.9 |
| Model Building and Refinement | | | | | |
| Initial models used | PDB:5ESV | PDB:4TOY | PDB:5ESV | | |
| | PDB:4TOY | PDB: 9NBT | PDB:4TOY | | |
| | PDB: 9NBT | | PDB: 9NBT | | |
| Model Composition | | | | | |
| Protein Chains | 12 | 10 | 12 | | |
| Protein Residues | 2407 | 2166 | 2407 | | |
| Ligands | 108 | 110 | 125 | | |
| Deviations from ideal (RMSD) | | | | | |
| Bond lengths outlier (#) | 0.012 (0) | 0.012 (0) | 0.011 (0) | | |
| Bond angles outlier (#) | 1.147 (12) | 1.204 (6) | 1.153 (32) | | |
| Ramachandran Plot | | | | | |
| Favored (%) | 99.28 | 99.53 | 98.88 | | |
| Allowed (%) | 0.72 | 0.47 | 1.12 | | |
| Outlier (%) | 0 | 0 | 0 | | |
| Validation | | | | | |
| Molprobity score | 0.69 | 0.81 | 0.7 | | |
| Clashscore | 0.57 | 1.09 | 0.61 | | |
| Poor rotamer (%) | 0.05 | 0.05 | 0.04 | | |
| EMRinger score | 3.6 | 3.74 | 2.17 | | |

**Table S1. Cryo-EM data collection, processing, and refinement validation statistics**
